# Mitophagy-related prohibitin 2 is an orthoflavivirus restriction factor targeted for degradation during infection

**DOI:** 10.64898/2025.12.05.692556

**Authors:** Imanol Rodrigo, María José Romero de Ávila, Belén García-Navarro, Martín Arellano, Laura Albentosa-González, Pilar Clemente-Casares, Edward Emmott, Armando Arias

**Affiliations:** Unidad de Medicina Molecular, Instituto de Biomedicina de UCLM (IB-UCLM), Universidad de Castilla-La Mancha (UCLM), Albacete, Spain; Unidad de Biomedicina UCLM-CSIC, Albacete, Spain; Instituto de Investigación Sanitaria de Castilla-La Mancha (IDISCAM), Spain; Facultad de Farmacia, UCLM, Albacete, Spain; Centre for Proteome Research, Department of Biochemistry, Cell & Systems Biology, Institute of Systems, Molecular & Integrative Biology, Biosciences Building, Crown Street, University of Liverpool, Liverpool, L69 7ZB, United Kingdom; Escuela Técnica Superior de Ingenieros Agrónomos y de Montes y Biotecnología (ETSIAMB), UCLM, Albacete, Spain

**Keywords:** Zika, Usutu, host-virus interactions, NS5 polymerase, NS2B/NS3 protease, myoferlin, restriction factors, proviral factors

## Abstract

1.

Orthoflaviviruses (OFVs) are the primary cause of arboviral disease worldwide, leading to large numbers of hospitalisations, sequelae and deaths associated. Viral genome replication is catalysed by non-structural protein 5 (NS5), the largest and most conserved protein in these viruses. Here, we used quantitative proteomics to identify host cell interactors of Zika virus (ZIKV), Usutu virus (USUV), and West Nile virus (WNV) NS5 proteins. A total of 141 proteins were detected in the analysis, being 26 of them associated with all three NS5s. Downregulation of myoferlin and LGALS3BP led to decreases in the virus yields, indicating that their presence is needed for efficient virus replication. Conversely, silencing of mitophagy-related prohibitin 2 (PHB2) resulted in significant increases in ZIKV and USUV titres and genomic RNA. Supporting a connection between this antiviral behaviour and mitophagy, an inhibitor of this pathway also led to larger virus titres and viral RNA copies. During infection, PHB2 protein levels gradually decreased, becoming transiently undetectable, and suggesting that it is specifically targeted by the virus for degradation. Ectopic expression of NS5 alone did not affect PHB2 intracellular abundance, however a significant decrease was observed in cells expressing the viral protease NS2B/NS3. Its elimination, was more pronounced in cells where both viral enzymes were co-expressed, supporting that NS5 could be assisting PHB2 recognition by NS2B/NS3. Overall, our results support that PHB2 is an orthoflaviviral restriction factor which is targeted for degradation to favour viral replication.

**IMPORTANCE:** Orthoflaviviruses (OFVs) are one of the main causes of disease transmitted by mosquitoes, with millions of infections occurring annually, and large numbers of deaths associated. Some important members include viruses causing dengue, yellow fever, and Zika-associated disease (e.g. microcephaly). OFVs contain a genome which is copied by a highly conserved viral protein termed NS5. Prohibitin 2 (PHB2) is a host cellular protein involved in different cellular processes including mitophagy, a mechanism to remove damaged mitochondria. Here, we show that human PHB2 interacts with orthoflaviviral NS5. Depletion of PHB2 results in increased viral yields, suggesting that this protein restricts orthoflaviviral infection. We found that PHB2 is targeted for degradation by the orthoflaviviral protease in a process aided by NS5, supporting that several viral proteins act concertedly to eliminate PHB2, counteracting its antiviral activity, and thus favouring infection.

## 3. INTRODUCTION

Over 50 per cent of the entire human population is at risk of infection by members of the *Orthoflavivirus* genus (1–3). Global warming and a broader geographic dissemination of competent vectors are factors contributing to a steady increase in the incidence and expansion of orthoflaviviral disease (4). Most human orthoflaviviruses (OFVs) are transmitted by mosquitoes, with estimates suggesting ∼100 million infections occur annually, leading to over 100,000 associated deaths (4, 5). The economic impact caused by OFVs is sizeable, with data supporting that Zika virus (ZIKV) and dengue virus (DENV) infections combined lead to >3 billion USD losses annually (6). ZIKV was declared a human pathogen of international concern in 2016, following large epidemic outbreaks that primarily affected the Americas. Now, the virus is present in six continents and more than 90 countries (7–9). Sequelae associated with infection during pregnancy can lead to severe congenital malformations, being microcephaly the most common syndrome (10). Although generally asymptomatic, infection in adults can cause clinical manifestations of different degree of severity, including meningitis, encephalitis and the Guillain-Barré syndrome (11, 12).

OFVs infecting birds as their natural vertebrate hosts are an additional source of public health concern (13). Even though humans are typically dead-end hosts for avian OFVs, neuroinvasive infection occur in a significant number of cases. West Nile virus (WNV) is the most relevant avian OFV from a public health perspective, with thousands of cases of severe disease and hundreds of fatalities annually (14). Despite that it generally leads to milder outcomes, a gradual increase in the number of neurological disorders associated with Usutu virus (USUV) has also been documented (15–18). WNV and USUV are genetically similar and exhibit serological cross-reactivity, which compromises their correct diagnosis in areas where cocirculation occur. Some lines of evidence imply that USUV infection in humans has been historically misdiagnosed as WNV, and its true impact underestimated (17, 19).

The OFV genome is a positive-stranded RNA molecule which is directly translated into a single polyprotein. This polypeptide is proteolytically processed by both cellular activities and the viral NS2B/NS3 protease, resulting in the release of different structural and non-structural proteins (20, 21). Non-structural protein 5 (NS5) contains a C-terminal RNA-dependent RNA polymerase (RdRp) domain, for the synthesis of new viral genomes, and an N-terminal methyltransferase (MTase) domain for the 5’ end capping of the viral genome (22). OFV replication factories are assembled inside partially secluded invaginations of endoplasmic reticulum (ER) membranes where viral RNA is isolated from host antiviral sensors while allowing the recruitment of replication factors (23–25). In addition to viral proteins, forty-six cellular proteins have been identified inside these vesicles, most of them displaying proviral activities (23).

The use of drugs targeting host factors required for viral replication has gained momentum in the antiviral research field (26, 27). In addition to viral replication, NS5 has been revealed as a multifaceted protein, engaging in diverse processes that include suppression of innate immunity, transcription reprogramming, and interference with the cell cycle progression (28–30). Several NS5 cellular partners have been identified, including proviral (23, 27, 31–34) and antiviral activities (35–37). Recent data have revealed that ZIKV NS5 binds to host protein Ajuba, preventing mitophagy activation (38). Mitophagy is a type of selective autophagy to remove irreparably damaged or depolarised mitochondria and promote cell survival. Since damaged mitochondria can activate innate immunity and proinflammatory responses, many viruses modulate the activity of this pathway (39). Ajuba knockdown *in vivo* resulted in enhanced inflammatory responses, leading to larger ZIKV dissemination in mouse tissues (38). These results convey that mitophagy suppression by NS5 favours viral spread in the host.

During mitophagy activation, PINK1 is accumulated in the outer mitochondrial membrane (OMM), leading to the recruitment of E3-ubiquitin ligase Parkin (40). Parkin ubiquitylates different OMM proteins, attracting other factors needed to initiate mitophagy (41). Prohibitin 2 (PHB2), a multitasking protein found in several subcellular locations, has been recently identified as a mitophagy receptor (40, 42). PHB2 interacts with SQSTM1 and LC3 in the inner mitochondrial membrane to drive damaged mitochondria to the autophagosome (43). Besides mitophagy, mitochondrial PHB2 in complex with prohibitin 1 (PHB1) has been proposed as major contributor to the MAVS-mediated antiviral signal, leading to the activation of type I interferon (IFN-I) responses (44). PHB2 is also found in the nucleus where it acts as a repressor of diverse transcription factors such as the oestrogen receptor (ESR), MyoD and MEF2 (45–48).

In the present study, we identified 141 host factors that interact with the NS5 proteins of ZIKV, WNV, and USUV. We found that 26 of these factors were common interactors for all three NS5s, including PHB2. PHB2 gene silencing led to increases in both virus titres and genomic RNA, which is suggestive of a viral restriction activity. During both ZIKV and USUV infection, PHB2 abundance gradually decreased, becoming transiently undetectable, and supporting that OFVs specifically target PHB2 to promote viral replication. Ectopic expression of NS2B/NS3 protease but not NS5 led to PHB2 degradation, although its elimination was more efficient when both viral proteins were co-expressed. In the study, we identified other NS5 interactors that are necessary for efficient virus infection, such as myoferlin (MYOF) and LGALS3BP, encouraging the use of specific drugs against them.

## 4. MATERIALS AND METHODS

### 4.1. Cells, viruses, drugs and cell viability assays

The origin of ZIKV and USUV strains, and protocols for infection have been detailed in previous manuscripts (32, 49–51). Cell culture infections and virus titrations by 50% tissue culture infectious dose (TCID_50_) assays have been performed following protocols described before (31, 32, 49). The antiviral efficacy of different host-targeting drugs has been tested as described in previous manuscripts (32, 51). To these assays, WJ460 [3-(3-(2-methoxyphenyl)-4-oxo-2-thiazolidinyl)-N-(4-phenylbutyl) benzamide] and mdivi-1 [3-(2,4-dichloro-5-methoxyphenyl)-2-mercaptoquinazolin-4(3H)-one] were purchased from MedChemExpress. The number of metabolically live cells after exposure to these compounds relative to mock-treated cultures, or after transfection with different host-targeting DsiRNA molecules relative to negative control DsiRNAs (not recognising human sequences) were obtained with the *CellTiterBlue* kit (Promega), as described before (31, 32).

### 4.2. Gene silencing with Dicer-substrate small-interfering RNAs (DsiRNAs)

Host-specific DsiRNA duplexes were obtained from Integrated DNA Technologies (IDT), with Information regarding their target sequences provided as supplemental material (Table S1). As a negative control, a non-specific DsiRNA (no target in the human genome), namely NonSP, was used. For gene silencing, 5 pmol of each DsiRNA were incubated during 20 min at room temperature in the presence of 0.25 μl of lipofectamine RNAiMAX (Invitrogen) and 50 µl of opti-MEM (Gibco). The mix was transferred into 96-plate wells, and then 1.5 x 10^4^ A549 cells were added on top. Twenty-four hours after transfection, the supernatants were removed, and virus was inoculated at an MOI of 0.1 or 1 TCID_50_/cell. Following adsorption, the inoculum was removed, cells washed, and fresh media containing 1% foetal bovine serum (FBS) added.

### 4.3. Viral RNA extraction and quantitative RT-PCR (RT-qPCR)

RNA was extracted from whole cell monolayers with the *NZY Total RNA Isolation* kit (NZYtech). ZIKV RNA was detected with a FAM-TAMRA probe targeting the E gene (1214 to 1244), and primers spanning positions 1192 to 1208 (forward) and 1269 to 1246 (reverse), while USUV RNA was detected using a FAM-TAMRA probe targeting the NS1 gene (3322 to 3345), and sense and antisense primers spanning positions 3288 to 3309 and 3372 to 3351, respectively. To the amplification of viral RNA, we used the *NZYSupreme One-step RT-qPCR Probe Master Mix (2x), ROX* kit (NZYtech), and RT-PCR conditions suggested by the manufacturer (15 min at 50°C, 3 min at 95°C, followed by 40 cycles of 5s at 95°C and 45s at 60°C). To obtain a standard curve with known amounts of RNA, we produced *in vitro* transcripts of the target genes. Briefly, 4μl of whole RNA extracted from infected cells were reverse transcribed using *SuperScript III Reverse Transcriptase* (Invitrogen). Then, 3μl of cDNA were PCR-amplified using *Phire Hot Start II DNA Polymerase* (ThermoFisher), and primers spanning residues 949 to 967 (forward) and 1934 to 1915 (antisense) for ZIKV, and 2809 to 2829 (forward) and 3808 to 3787 (antisense) for USUV (both forward primers contained a T7 promoter. PCR amplicons were purified (*NZY GelPure* kit, NZYtech), and ligated into the PCR Blunt plasmid (*Zero Blunt PCR cloning* kit, ThermoFisher), and the linearised plasmids or directly the PCR amplicons were *in vitro* transcribed with the *mMESSAGE mMACHINE T7 Transcription* kit (Ambion). After DNAse treatment, the RNA transcript was purified and quantitated, and the standards were prepared with known amounts of each transcript in 100 ng/μl yeast RNA.

### 4.4. Relative quantification of host mRNAs and poly I:C treatment

To confirm host gene knockdown after DsiRNA transfection, host gene mRNA levels were determined with *One-step NZYSpeedy RT-qPCR Green kit, ROX* kit (NZYtech) in a CFX Duet Real Time PCR instrument (BioRad) and specific primer pairs, all provided by IDT [reference numbers (Hs.PT.58.): 27059869, 19355804, 40826680, 2219540, 3098270, 40209489, 456578.g, 3838519.g, 38465792, 22859503, 22973334, 27422678.g, 40506149, 40261049]. As a a housekeeping gene we used primers amplifying the GAPDH transcripts (Hs.PT.39a.22214836). The efficiency of downregulation for each transcript was calculated following the 2^-ΔΔCT^ method (32, 52).

To stimulate IFN-I expression, A549 cells were transfected with *Poly (I:C) (LMW)* (Invivogen), using Lipofectamine 3000 in the absence of the P3000 reagent (Invitrogen). Twenty-four hours prior poly I:C stimulation, cells were transfected with DsiRNAs (either targeting PHB2 or a non-specific DsiRNA), as explained above. Total RNA was extracted from poly I:C-stimulated cells to analyse IFN-β gene transcription (relative IFNB1 mRNA; a primer pair purchased from IDT, reference Hs.PT.58.39481063.g).

### 4.5. Vectors and mutagenesis

Constructs expressing HA-tagged versions of full-length NS5 (HA-NS5) or the RdRp domain alone (HA-RdRp) from ZIKV, USUV, and WNV were used as documented in previous manuscripts (31, 32, 53). To obtain the ZIKV NS2B/NS3 protease vector (pZIKV-NS2B3), we used a pcDNA3 clone expressing ZIKV RdRp fused to an HA tag and the EYFP (pZIKV-HA-RdRp-EYFP) as a backbone. We swapped the insert in this vector by the corresponding NS2B3-coding region, using the *In-Fusion HD EcoDry Cloning Kit* (Takara Bio), using specific reverse (5’-AGCGTAATCTGGAACATCGTATGGGTA-3’) and forward (5’-GATAACTCGAGCATGCATCTAGAG-3’) primers, leading to the amplification of the vector devoid of the RdRp-EYFP insert. To the amplification of the NS2B3 coding region we used ZIKV primers spanning genomic positions 4224 to 4244 (forward) and 6464 to 6444 (antisense). Both primers contained 15-nt tails overlapping with the linear vector, enabling the ligation of the insert by recombination.

### 4.6. Co-immunoprecipitation and mass spectrometry

For coimmunoprecipitation assays, HA-NS5 or HA-RdRp plasmids were transfected in A549 or 293T cells with either the *Lipofectamine 3000* kit (Invitrogen) or the *FuGENE 4K* reagent (Promega). At 24 to 48 h post-transfection, cells were scrapped on ice-cold PBS, washed in PBS twice, spun at 300x *g* for 5 min, and the pellet resuspended at 4°C in 450 μl of RIPA buffer (10 mM Tris-HCl pH 7.5; 150 mM NaCl; 0.5 mM EDTA; 1% SDS; 1% Triton X-100; 1% deoxycholate) in the presence of 1x *Complete Mini EDTA-free protease inhibitor cocktail* (Merck) and 250 U/ml benzonase (Sigma). To remove the cellular debris, the lysate was spun during 10 mins at 15000x *g*. Protein concentration in the supernatants was determined with the *BCA Protein Assay Kit* (Pierce), and all samples normalised to the same concentration in a final volume of 400 μl. The samples were then equilibrated in 1.5 volumes (600 μl) of dilution buffer (10 mM Tris-HCl pH 7.5; 150 mM NaCl; 0.5 mM EDTA), and then mixed with either agarose or magnetic anti-HA beads (Pierce), and incubated in a rotary shaker overnight at 4°C. The beads were then either magnetically separated (magnetic) or pelleted by centrifugation (agarose), the supernatant eliminated, and washed four times in 1x dilution buffer containing 0.05% Nonidet P40 substitute (Fluka BioChemika). Finally, the co-immunoprecipitated proteins were released from the beads by heating at 90°C for 5 min in 1x *Bolt LDS Sample Buffer* (Life Technologies).

For the identification of host cell interactors by mass spectrometry, HA-NS5 proteins expressed in A549 cells were pulled down with anti-HA magnetic beads, as described above. The pulldowns were SP3-precipitated (54), trypsin digested and labelled with TMTpro. LC-MS/MS was performed on a Dionex U3000 UHPLC, in-line with a Q-Exactive-HF mass spectrometer as described previously (55). Data were analysed with MaxQuant 1.6.7.0 (56). N-terminal protein acetylation and methionine were set as variable modifications, enzyme specificity was set to trypsin with up to two missed cleavages. Data were searched against fasta files containing the Swissprot human proteome entries (20,300 entries, downloaded 2020/03/12) and a custom fasta file containing the HA-tagged NS5 sequences. For each NS5, four independent transfections and co-IPs were performed, one control was removed at the data analysis stage as an outlier (Fig. S1). To identify those factors that were significantly enriched in the IPs, four independent plates were mock-transfected in parallel with an empty vector (pcDNA3). NS5-interacting factors were identified using the *Perseus software* version 2.0.6.0., as previously described (57). A list with all host factors that are significantly enriched in ZIKV, USUV and WNV HA-NS5 co-IPs followed by TMT-based proteomics are provided as supporting material (Tables S2 – S4). The mass spectrometry data, including .fasta files has been submitted to proteomeXchange via the PRIDE repository and has accession number PXD071636.

### 4.7. Western blot analysis

To the detection of host and virus proteins, we followed protocols available in previous studies (32). Antibodies against the HA epitope (BioLegend, reference 901501), the FLAG epitope (Proteintech, reference 20543-1-AP), PHB2 (Invitrogen, PA5-21417), STAT2 (Cell Signaling, 72604S), ZIKV NS3 (Invitrogen, PA5-143436) and ZIKV NS5 (Genetex, GTX133328) were used. As loading controls, we purchased HRP-conjugated antibodies against GAPDH (Genetex, GTX627408-01) and tubulin (Cell Signaling, 12351S).

### 4.8. Gene ontology analysis

Gene Ontology (GO) enrichment analysis was performed on differentially abundant proteins (FDR < 0.05) using g:Profiler with the Benjamini-Hochberg correction. Redundant GO Biological Process (BP) terms were reduced using rrvgo (v1.10.0) with a Wang semantic similarity threshold of 0.7. GO Cellular Component (CC) terms with <5 genes were excluded. Remaining terms were grouped only when sub-compartments shared a parent GO category (e.g., mitochondrial compartments) and the combined gene count remained statistically robust (≥5 genes). A summary of all enrichment analysis to the identification of BP and CC terms are provided as supporting information (Tables S5 and S6).

## 5. RESULTS

### 5.1. Identification of host factors associated with orthoflaviviral NS5s

Full-length versions of NS5 proteins were expressed in A549 cells, and their interacting host factors isolated as described in the Methods section. Mass spectrometry analyses resulted in the identification of 72, 63 and 77 cellular proteins significantly enriched in pulldowns of ZIKV, USUV and WNV NS5s, respectively (Fig. 1, Tables S1 – S4). The GO Biological Process enrichment analysis revealed that most of them were involved in RNA metabolism (splicing and processing), ribosome biogenesis and protein translation, chromatin assembly and chromosome organisation, or respiration and proton transport in the mitochondria (Fig. 1 and 2, and Fig. S2). The intersection of the three sets revealed that 26 proteins were common interactors for ZIKV, WNV and USUV NS5s (Fig. 1A and Fig. S2). To the identification of those factors that could be more biologically relevant, we re-examined the intersection, but only using the top 50 significantly enriched proteins in each pulldown. This resulted in 14 host factors shared by all three sets (Fig. 2, Tables S2 – S4). Most of them are mitochondrial proteins (ATP5B, PHB2, PGAM5, FH and SFXN1) or participate in ribosome biogenesis and/or protein translation (RSL1D1, RPL21, RPL24, RPL24).

**Fig. 1.**
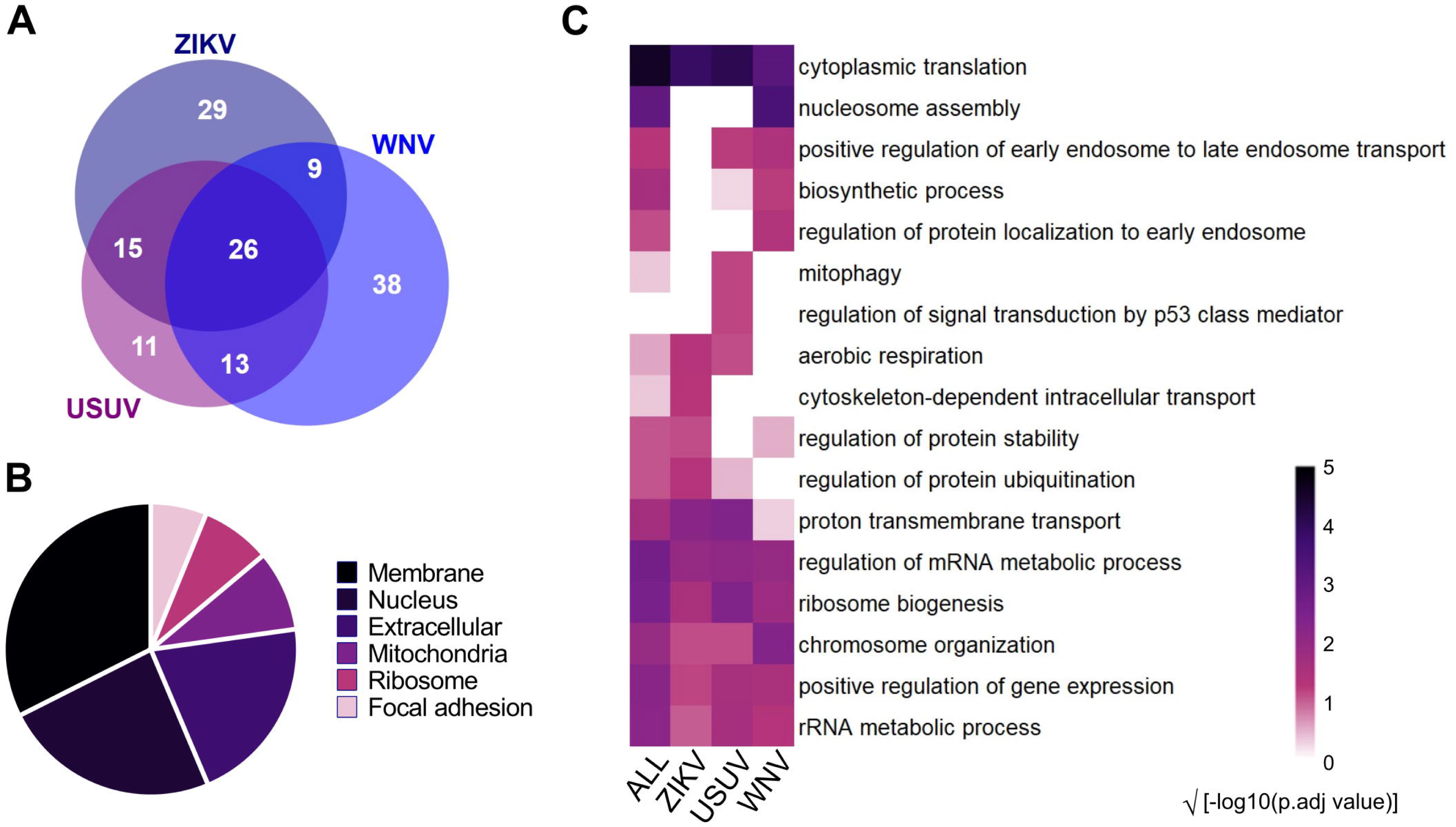
Summary of orthoflaviviral NS5-interacting network. (A) Overlap of host proteins interacting with different orthoflaviviral NS5 proteins (ZIKV, USUV and WNV). BioVenn was used to the graphic representation of the diagram (85). (B) Gene Ontology analysis of cellular components (GO-CC). Related sub-compartments, i.e. mitochondrial inner membrane (GO:0005743) and mitochondrial nucleoid (GO:0042645) were grouped as ’Mitochondria’ to reflect organelle-level localization. The pie chart displays the proportional distribution of significantly enriched terms (adjusted p < 0.05) among all factors identified in NS5-interacting datasets. (C) Heatmap of significantly enriched Gene Ontology Biological Process (GO-BP) terms in co-IPs of cells transfected with either ZIKV, USUV, or WNV HA-tagged NS5 proteins (adjusted *p* < 0.05). Each GO-BP is represented in a colour-based gradient for statistical significance [√(−log_10_ adjusted p-value]. GO terms were clustered hierarchically using semantic similarity (rrvgo; similarity threshold = 0.7). See Tables S5 and S6 for full terms.

**Fig. 2.**
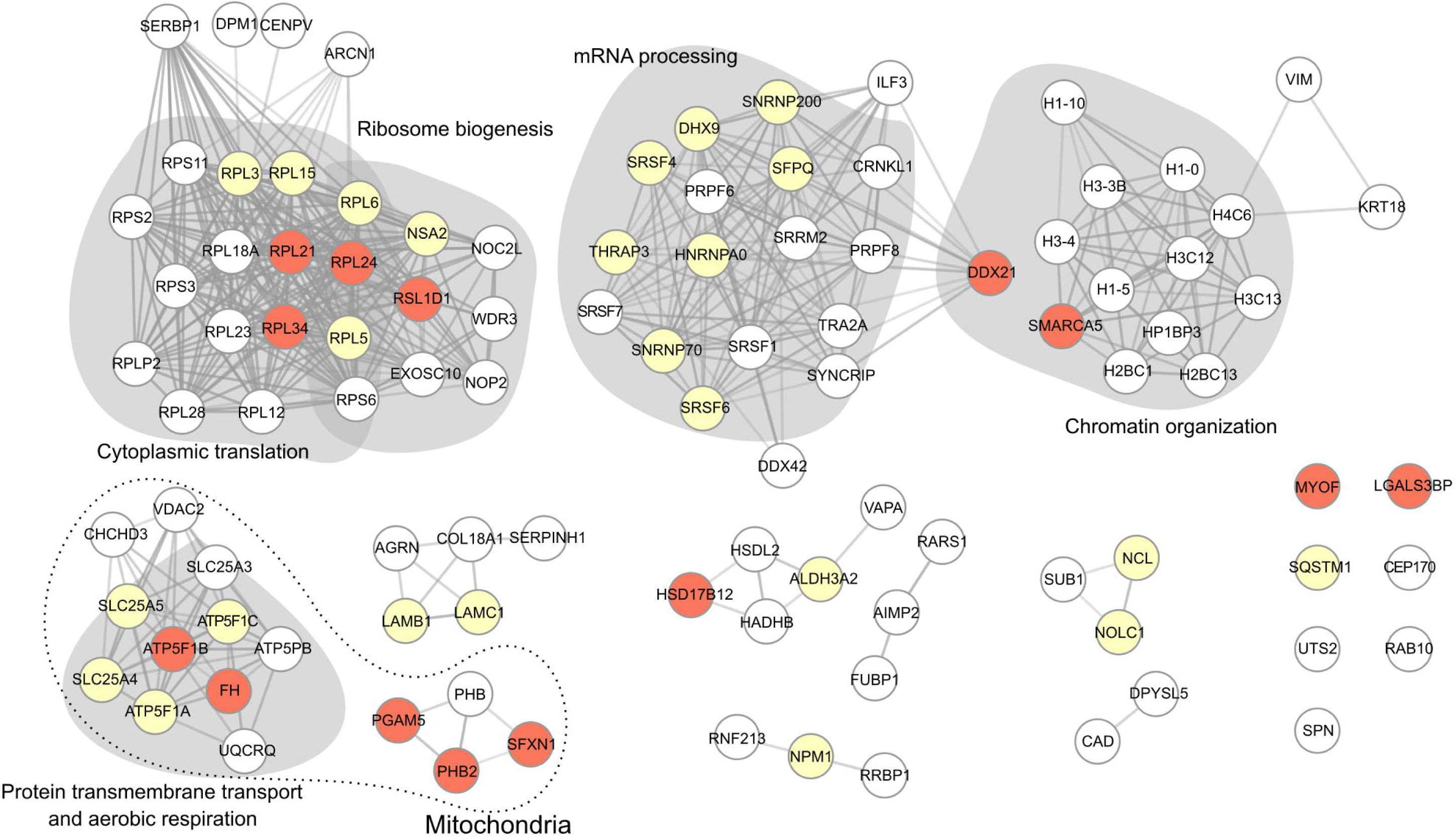
Orthoflavivirus NS5 interaction network. Quantitative proteomics was used as described in the text to identify host factors interacting with HA-tagged versions of ZIKV, USUV or WNV NS5 proteins. Only the top 50 most significantly enriched factors in each data set are used to build the network and to identify overlaps between data sets. The protein-protein interaction network was generated using the STRING database (86). Data clustering was generated in Cystoscape (87), using the MCL algorithm. Those factors identified in two datasets are coloured in pale yellow, and those in all three datasets in coral. Each grey-shaded area represents a cluster positively associated with a biological process (i.e. ribosome biogenesis, cytoplasmic translation, mRNA processing, chromatin organisation, proton transmembrane transport/aerobic respiration). Mitochondrial proteins are encircled by a dotted line.

### 5.2. Silencing of NS5-host partners identifies proviral and restriction activities

To elucidate the biological impact of these 14 host factors to OFVs, we examined the effect of their silencing during infection (Fig. 3). All the DsiRNAs used in this study elicited efficient gene downregulation, with over 70% reduction in mRNA levels compared to controls (Fig. S3). Except for ribosomal proteins RPL21, RPL24 and RPL34, gene silencing had no effect or caused only a modest decrease in cell viability (>75% live cells) at 48 h post-transfection (Fig. S4). Downregulation of either PHB2 or ribosomal L1 domain-containing protein 1 (RSL1D1) resulted in larger virus titres in the supernatant of both ZIKV- and USUV-infected cells, suggesting these two proteins are restriction factors for the OFVs (Fig. 3). Downregulation of RPL21, PGAM5, SFXN1 and SMARCA5 also led to significant increases in USUV but not in ZIKV titres, while DDX21 silencing resulted in increased infectivity for ZIKV, but not for USUV.

**Fig. 3.**
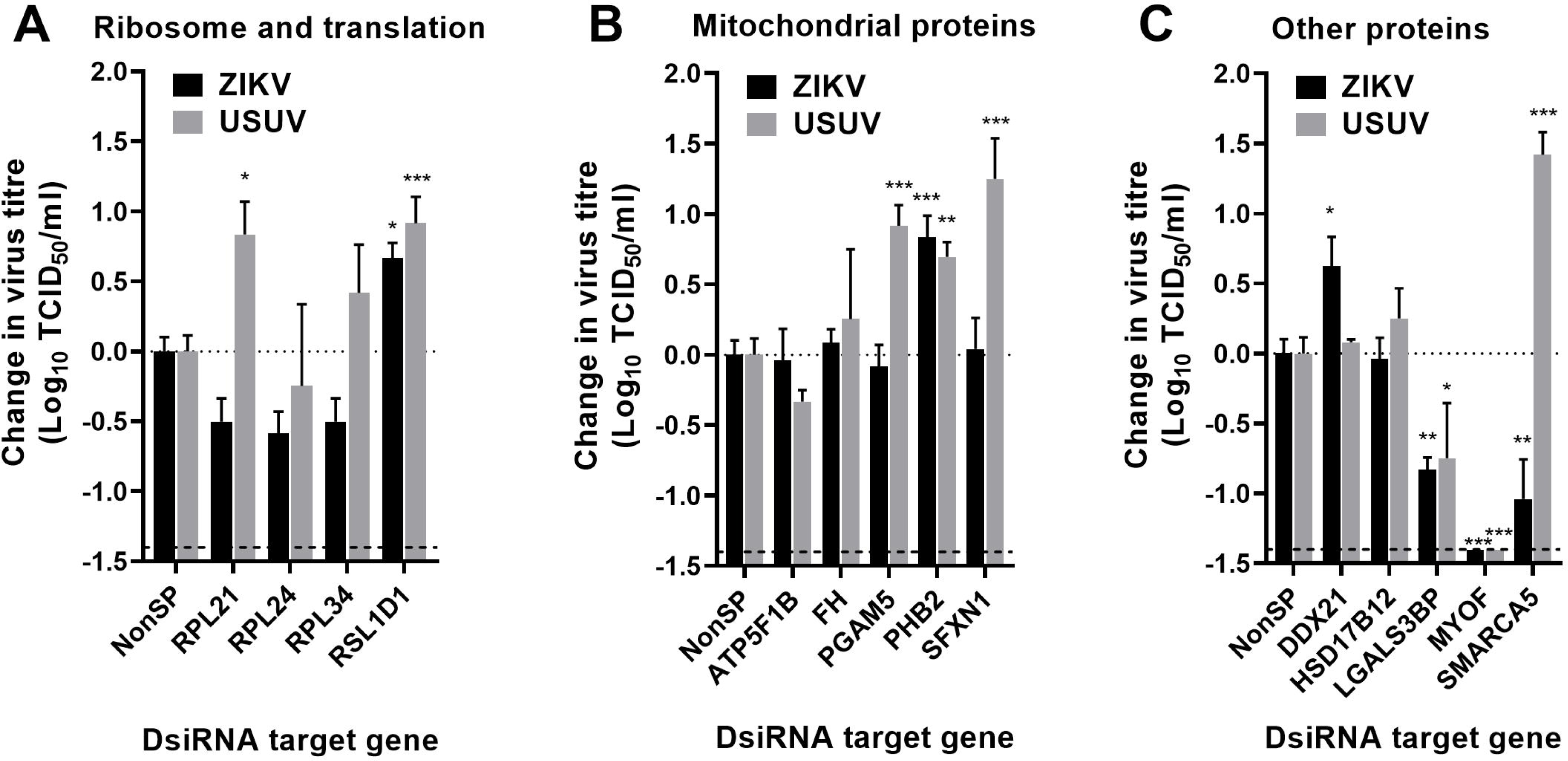
DsiRNA-mediated gene silencing identifies putative proviral and restriction host factors for OFV replication. Virus titres in the cellular supernatant of infected cells (n ≥ 4 biologically independent samples), previously transfected with a DsiRNA targeting an NS5-interacting factor. At 24 h post-transfection, A549 cells were infected with either ZIKV (black columns) or USUV (grey columns) at an MOI of 0.1 TCID_50_/cell. Control infections were carried out in parallel experiments in cells transfected with a non-human target DsiRNA (NonSP). The bars represent relative changes in the viral titres of gene-silenced cultures compared to NonSP controls, both at 48 hpi. To improve clarity, the effect of host factor downregulation on viral infectivity is represented in 3 different graphs: ribosome biogenesis and translation (A), mitochondrial (B), and other ungrouped host factors (C). Significant differences observed between unspecific (NonSP) and specific host-targeting DsiRNA-transfected cells are indicated (Two-way ANOVA, Dunnett’s multiple comparisons test; *, p < 0.05; **, p < 0.01; ***, p < 0.001). Bars are representing the standard error of the mean (SEM) for each data point.

Silencing of LGALS3BP and MYOF negatively affected both ZIKV and USUV, with pronounced losses observed in the virus titres (Fig. 3). Additional analyses at different time points and MOIs further confirmed this observation, conveying that these two are proviral proteins (Fig. 4 and Fig. S5). Viral replication was significantly impaired when MYOF was silenced, with 1,000- to 10,000-fold decreases in virus titres at 72 h post-infection (hpi). These findings enticed us to evaluate whether specific drugs against MYOF, such as WJ460, exhibit an antiviral behaviour. Although WJ460 specifically inhibits all known MYOF cellular functions (58), we observed only a modest reduction in ZIKV titres, whilst no effect on USUV was found (Fig. 4E – 4G). This apparent lack of antiviral effect was observed even at concentrations causing significant cell toxicity (40 to 160 μM), in contrast to data obtained with DsiRNAs.

**Fig. 4.**
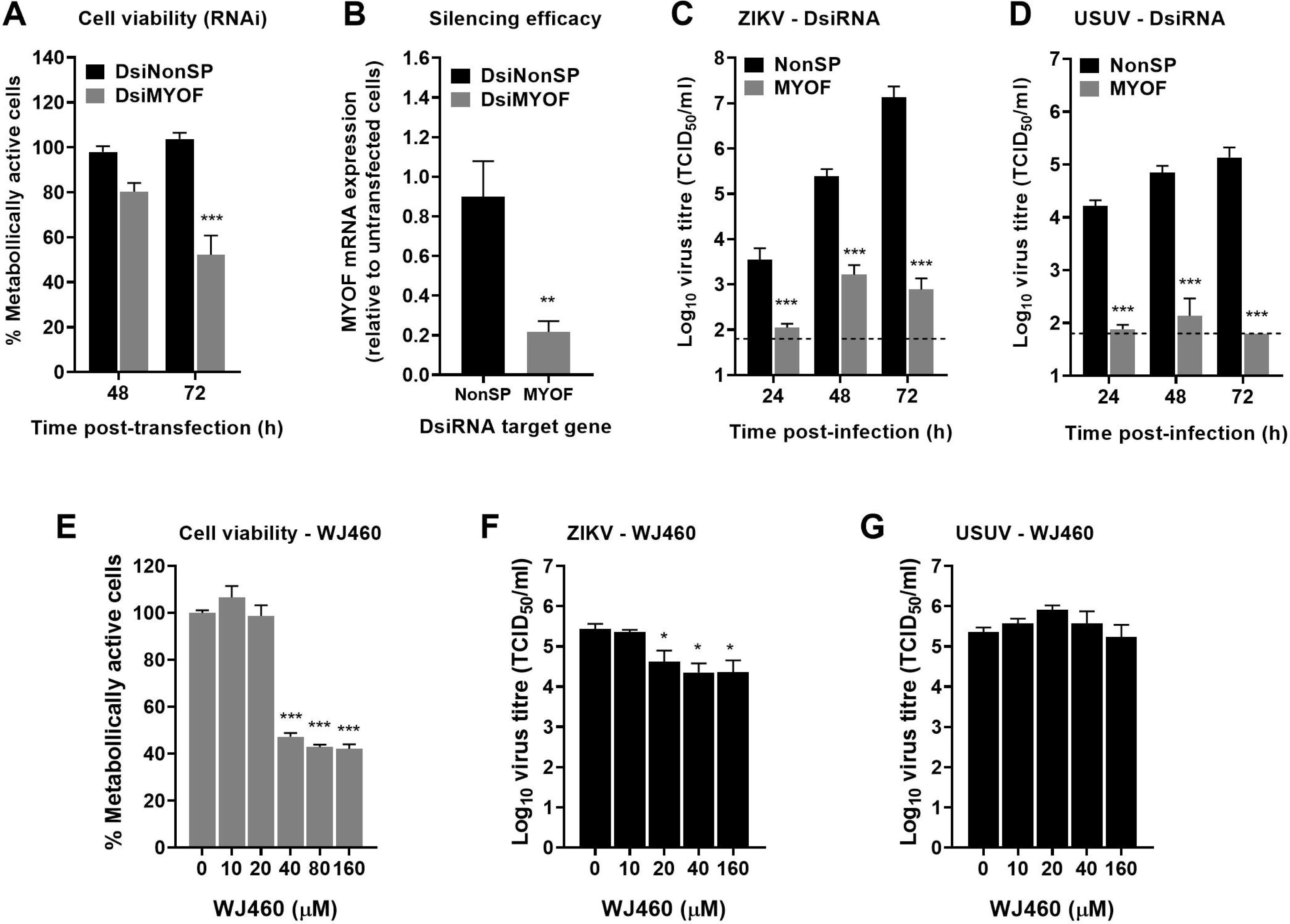
Reduced virus titres in MYOF-depleted cells. (A – D) Effect of MYOF gene silencing on viral replication. (A) Relative amount of metabolically active cells after transfection with a specific (DsiMYOF) or an unspecific DsiRNA (DsiNonSP). Cell viability is determined at 48 and 72 h post-transfection as suggested in the Methods section. Values are obtained as a percentage relative to mock untransfected cells. (B) Relative MYOF mRNA expression was examined by RT-qPCR in A549 cells transfected with either a MYOF-targeting (DsiMYOF) or a non-human target DsiRNA (DsiNonSP). MYOF mRNA levels were normalised to GAPDH mRNA levels. Values obtained are the average of at least 4 independent biological replicas (n ≥ 4), and relative to untransfected cells carried out in parallel. (C, D) Virus infectivity was examined by TCID_50_ assays of samples collected at different time points from cells previously transfected with each DsiRNA. Twenty-four hours after transfection, cells were infected with ZIKV (C) or USUV (D) at an MOI of 1 TCID_50_/cell. Each value in the graph is the average of ≥ 4 independent biological replicas. (E – G) Effect of MYOF-targeting drug WJ460 on ZIKV and USUV replication. (E) The relative amount of metabolically active cells after treatment during 48 h with increasing concentrations of WJ460 is calculated as indicated in (A). Values are represented as a percentage relative to mock-treated cells. (F, G) ZIKV (F) or USUV (G) titres detected in cells treated with WJ460 at 48 h postinfection. In each panel, significant differences are indicated (Two-way ANOVA, Sidak’s multiple comparisons test; *, p<0.05; ***, p < 0.001). The standard error of the mean (SEM) is represented for each data point.

### 5.3. PHB2 downregulation stimulates orthoflaviviral replication

Since the preliminary DsiRNA-based screening identified PHB2 and RSL1D1 as putative restriction factors (Fig. 3), we further examined how their silencing affected virus replication (Fig. 5 and Fig. S6). In agreement with our previous observations, downregulation of these genes resulted in significantly larger ZIKV and USUV titres at different time points and MOIs tested. Owing to precedent studies reporting that ZIKV NS5 interferes with mitophagy to promote viral dissemination (38), and that PHB2 is a mitophagy receptor, we aimed at further characterising this protein during the virus life cycle. In addition to enhanced virus titres, PHB2 silencing also led to significant increases in intracellular viral RNA (Figs 5G – 5J) that was also accompanied with increases in viral protein NS5 expression (Fig. 5B and Fig. S7), further supporting that PHB2 restricts viral replication. Treatment of infected cells with mdivi-1, an inhibitor of mitochondrial fission-related protein DRP1 which represses mitophagy (59), also resulted in both larger virus titres and genomic RNA (Fig. 6 and Fig. S8). Altogether, these results are pointing towards mitophagy and PHB2, a key receptor in this pathway, as restriction factors of the orthoflaviviral life cycle.

**Fig. 5.**
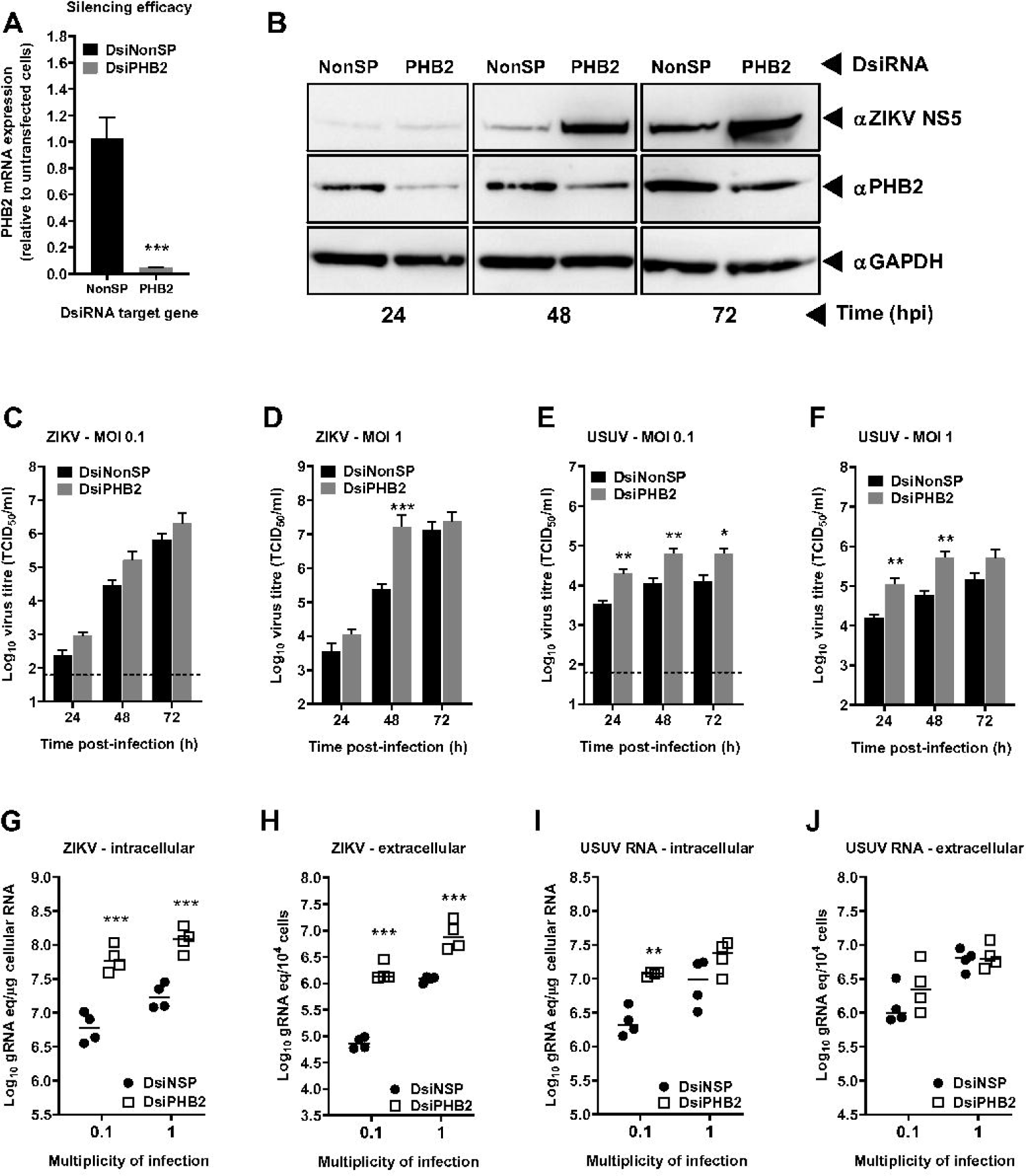
Increased viral titres, RNA and protein levels in PHB2-depleted cells. (A) Relative PHB2 mRNA expression in A549 cells previously transfected with either a PHB2-targeting (DsiPHB2) or a non-human target DsiRNA (DsiNonSP). PHB2 expression was normalised to GAPDH mRNA levels. Values obtained are the average of at least 4 independent biological replicas (n ≥ 4), and relative to untransfected cells carried out in parallel. (B) PHB2 downregulation leads to increased ZIKV NS5 protein abundance in infected cells. Representative WB of ZIKV NS5 and PHB2 protein levels in A549 cells previously transfected with each DsiRNA and infected at 24 h post-transfection with ZIKV. As a loading reference, an anti-GAPDH antibody was used. (C – F) Virus infectivity was examined in the supernatants of cells previously transfected with each DsiRNA. Twenty-four hours after DsiRNA transfection, cells were infected with ZIKV (C, D) or USUV (E, F) at an MOI of 0.1 (C, E) or 1 TCID_50_/cell (D, F). Each value in the graph is the average of four independent biological replicas (n = 4). (G – J) Viral RNA genome equivalents detected in samples collected at 48 hpi from cells previously transfected with each DsiRNA. Twenty-four hours after DsiRNA transfection, cells were infected at an MOI of 0.1 or 1 TCID_50_/cell with ZIKV (G, H) or USUV (I, J). Each value in the graph represents the average of viral RNA detected in four independent biological replicas (n = 4) extracted from infected cells (intracellular, graphs G and I) or the supernatant of these cultures (extracellular, graphs H and J). The statistical significance in each panel was determined by Two-way ANOVA and Sidak’s multiple comparison test (C – J) or *t* (A) tests (*, p<0.05; **, p<0.01; ***, p < 0.001). Bars are representing the standard error of the mean (SEM).

**Fig. 6.**
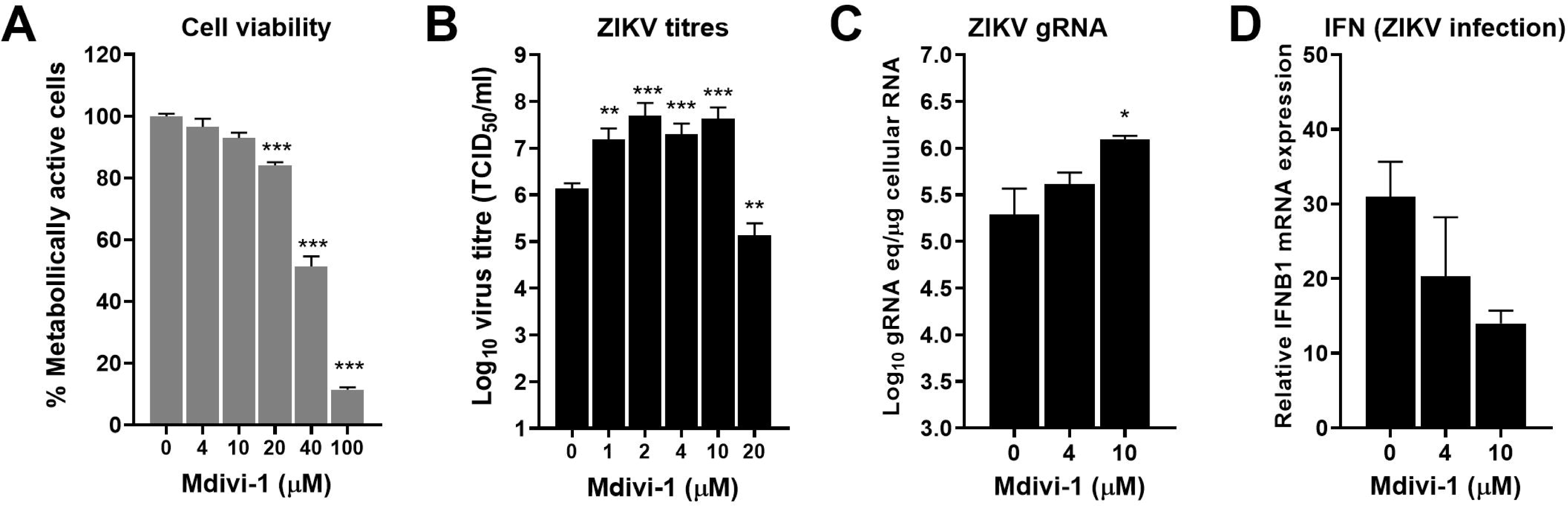
Increased ZIKV titres and RNA levels in cells treated with a mitophagy inhibitor. (A) Relative number of metabolically active A549 cells after 48 hours in the presence of increasing concentrations of mdivi-1. Values are relative to untreated cells (0 μM). (B) ZIKV titres at 48 hpi in the supernatant of cells treated with increasing concentrations of mdivi-1. Cells were previously inoculated with ZIKV at an MOI of 0.1 TCID_50_/cell. (C) Genomic RNA equivalents in cells infected with ZIKV, and treated with mdivi-1. D, Relative IFNB1 mRNA expression in ZIKV-infected cells treated with mdivi-1.

To examine whether larger titres and RNA levels could be related to an increase in virus particle release, we compared extracellular to intracellular viral RNA in PHB2-depleted and unspecific DsiRNA-treated cells (Fig. S9), but no significant differences were found. This observation imply that larger virus titres and genomic RNA in PHB2-depleted cells are likely the result of enhanced genome replication, and hence PHB2 is an OFV antagonist.

### 5.4. PHB2 is transiently eliminated during infection

Owing its apparent influence on viral replication, we decided to examine PHB2 protein abundance dynamics during infection (Fig. 7). We found that PHB2 intensity gradually decreased to become barely detectable at 48 hpi (< 3% relative to mock-infected cells) in both ZIKV- and USUV-infected A549 cells (Fig. 7). At later time points, PHB2 intensity was detected at a similar level than in control uninfected cells (101% and 58% at 60 hpi, in ZIKV- and USUV-infected cells, respectively). PHB2 signal was also transiently reduced in 293T cells, supporting that is either targeted for degradation or its expression repressed (Fig. 7H). Since mock- and virus-infected cells exhibit no significant differences in the amount of PHB2 mRNA (Fig. S10), reduced protein levels are possibly connected to degradation.

**Fig. 7.**
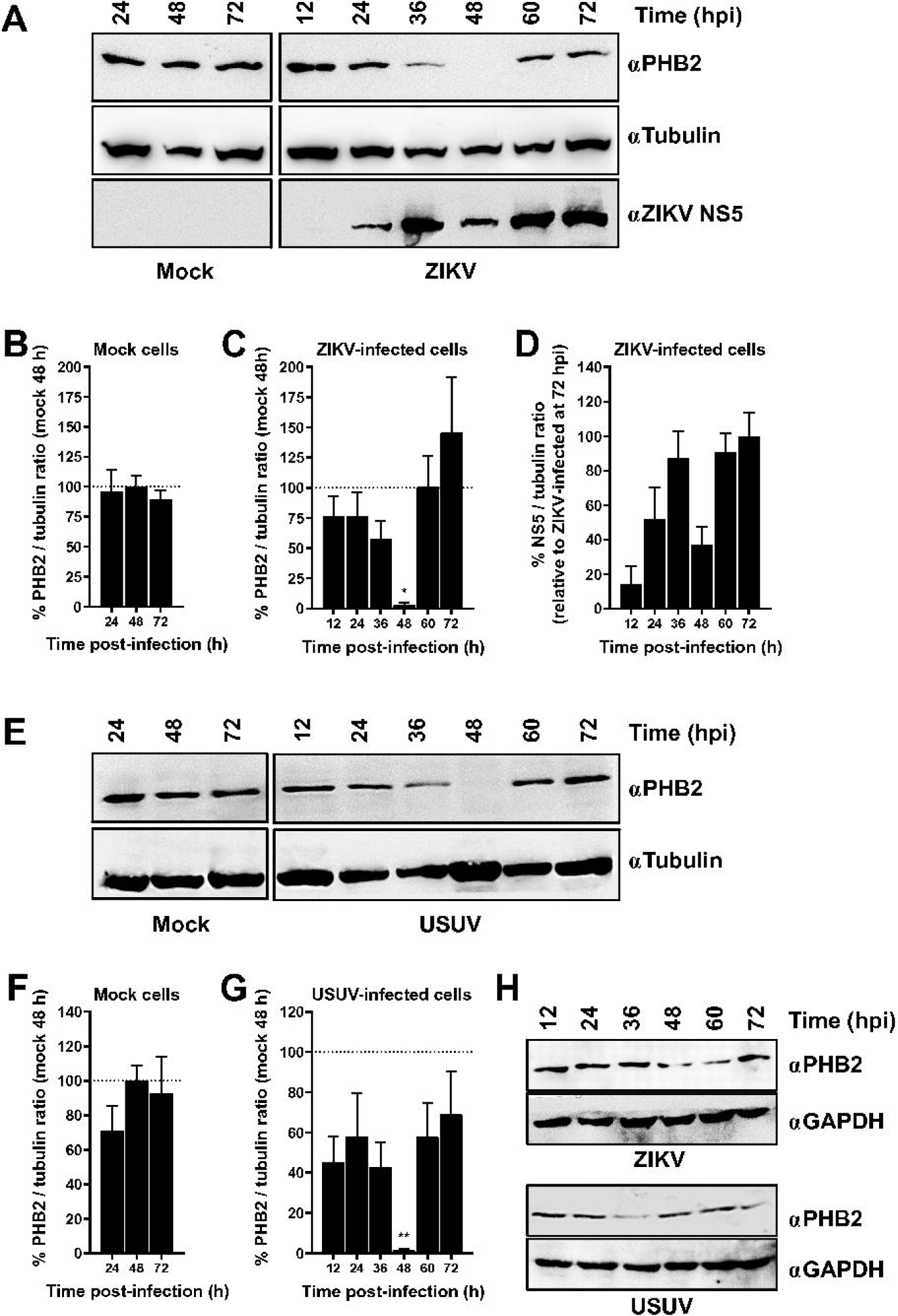
PHB2 is transiently eliminated in ZIKV- and USUV-infected cells. (A, E) Representative Western blot of PHB2 kinetics in cells infected with either ZIKV (A) or USUV (E). (B, C, F, G) PHB2 protein levels in mock- (B, F), ZIKV- (C) or USUV-infected (G) A549 cell lysates at different time points. PHB2 to tubulin band intensity ratios are relative to those obtained at 48 h in Mock-infected cultures. (D) ZIKV NS5 protein levels detected in ZIKV-infected A549 cell lysates collected at different time points. NS5 to tubulin band intensity ratios for each time point are relative to 72 hpi. Each value in the graphs is the average of three independent biological replicas (n = 3). Bars are representing the standard error of the mean (SEM) for each data point. Significant differences between mock and infected samples are indicated (One-way ANOVA; *, p < 0.05; **, p < 0.01). (H) Representative blot of PHB2 protein levels in 293T cells collected at different time post-infection in cells infected with ZIKV (top) or USUV (bottom). An anti-GAPDH antibody was used to calibrate total protein loading.

### 5.5. PHB2 depletion correlates with larger interferon expression in infected cells

Since PHB2 has been linked to MAVS-signalling, we considered that its degradation may be interfering with IFN-I expression. To assess this possibility, we analysed IFN-β gene (IFNB1) transcription in cells previously knocked down for PHB2. In both ZIKV- and USUV-infected cultures, we found larger IFNB1 mRNA levels when PHB2 was silenced than in controls transfected with a non-specific DsiRNA (Figs 8A – 8D), in contrast with previous observations (44). A reasonable scenario is that enhanced IFNB1 mRNA transcription could be incidental rather than directly prompted by PHB2 knockdown, given that more viral RNA genomes are detected in PHB2-depleted cells (Figs. 5G and 5I). Thus, we quantitated the relative amount of IFNB1 mRNA in cells stimulated with poly I:C, a non-replicating double-stranded RNA. Although less pronounced than in infected cells, PHB2 silencing also resulted in increased IFNB1 transcription (Figs 8E – 8H). To further explore whether IFN-I activation was connected to mitophagy inhibition, we analysed transcription levels in ZIKV-infected cells treated with mdivi-1, but in this case, IFNB1 mRNA levels were lower (Fig. 6D).

**Fig. 8.**
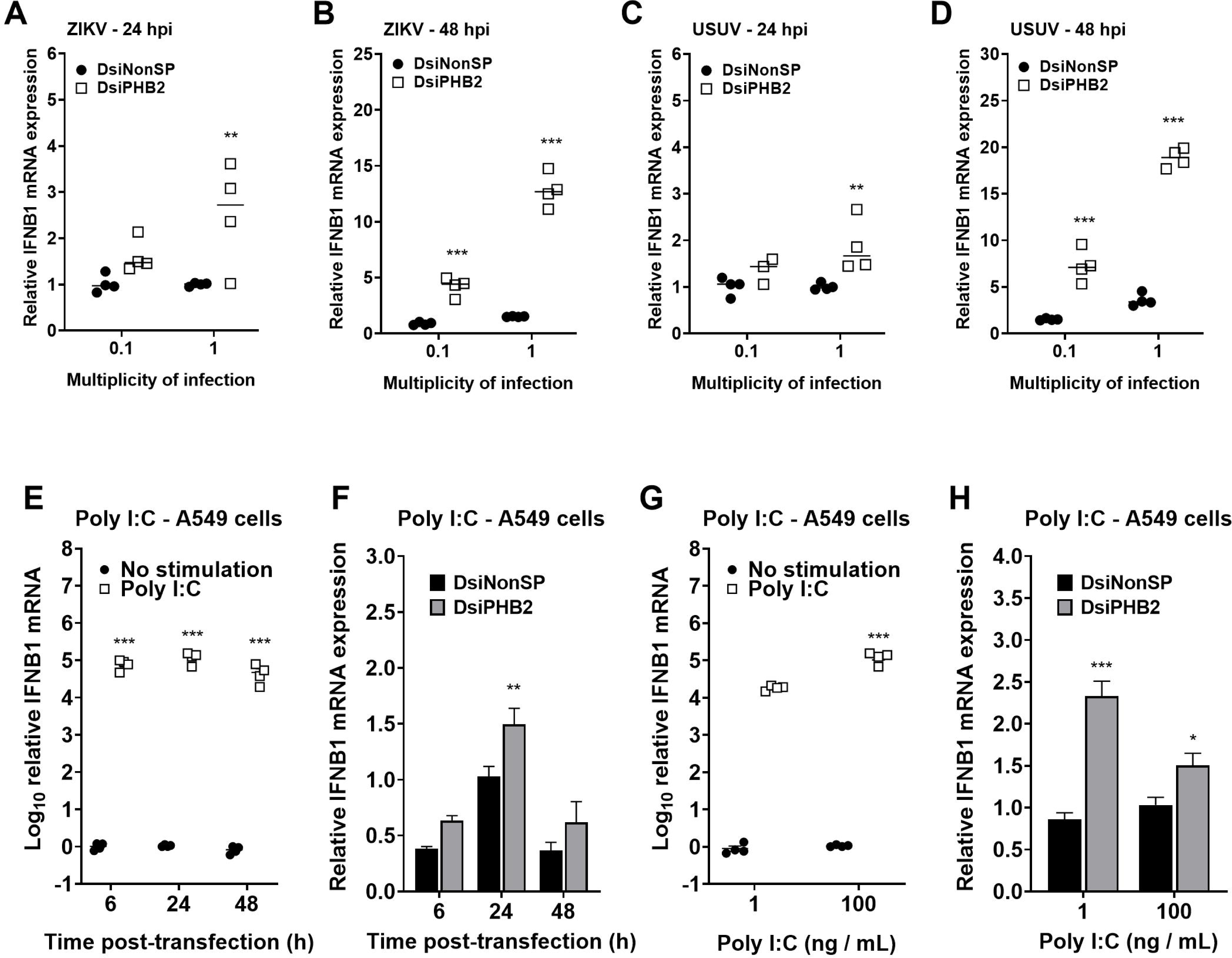
Increased IFN-I transcription in PHB2-downregulated cells. (A – D), Relative IFNB1 mRNA expression in A549 cells previously transfected with either a PHB2-targeting (DsiPHB2) or a non-human target (DsiNonSP) DsiRNA, and then infected with ZIKV (A, B) or USUV (C, D). IFNB1 expression was normalised to GAPDH mRNA levels. (E – H) Relative IFNB1 mRNA expression in A549 cells transfected with either a PHB2-targeting (DsiPHB2) or a non-human target (DsiNonSP) DsiRNA, and then stimulated with different amounts of poly I:C. IFNB1 expression was normalised to GAPDH mRNA levels. Values obtained are the average of at least 4 independent biological replicas, and relative to control cells (no DsiRNA). (E, G) IFNB1 mRNA levels in poly I:C-stimulated cells compared to control unstimulated cells. (F, H) Relative IFNB1 mRNA expression in PHB2-depleted cells compared to control cells (DsiNonSP) after poly I:C stimulation.

### 5.6. NS5 stimulates PHB2 degradation by the viral protease NS2B/NS3

Since both ZIKV and DENV NS5 directly promote host STAT2 degradation by the proteasome (35, 37), and that NS5 specifically interacts with PHB2 (Fig. 2), direct PHB2 degradation by NS5 was a plausible scenario. Further supporting this possibility, ZIKV NS5 and host PHB2 protein levels coincidentally decrease at 48 hpi in A549 cells when compared to earlier time points (Fig. 7). We first confirmed that PHB2 and NS5 interact in co-IP assays followed by Western blot (Fig. 9A), as inferred from mass spectrometry data (Tables S2 – S4). PHB2 was also pulled down when the RdRp alone was used as a bait, supporting that the interaction occurs through the polymerase domain (Fig. 9B). To assess whether NS5 expression and PHB2 degradation were linked, we used increasing amounts of all three NS5 constructs, but no significant change in PHB2 was detected (Fig. 9C and Fig. S11). As expected, the expression of recombinant ZIKV NS5 significantly affected STAT2 protein levels detected (Fig. 9C and Fig. S11), as previously documented (37).

**Fig. 9.**
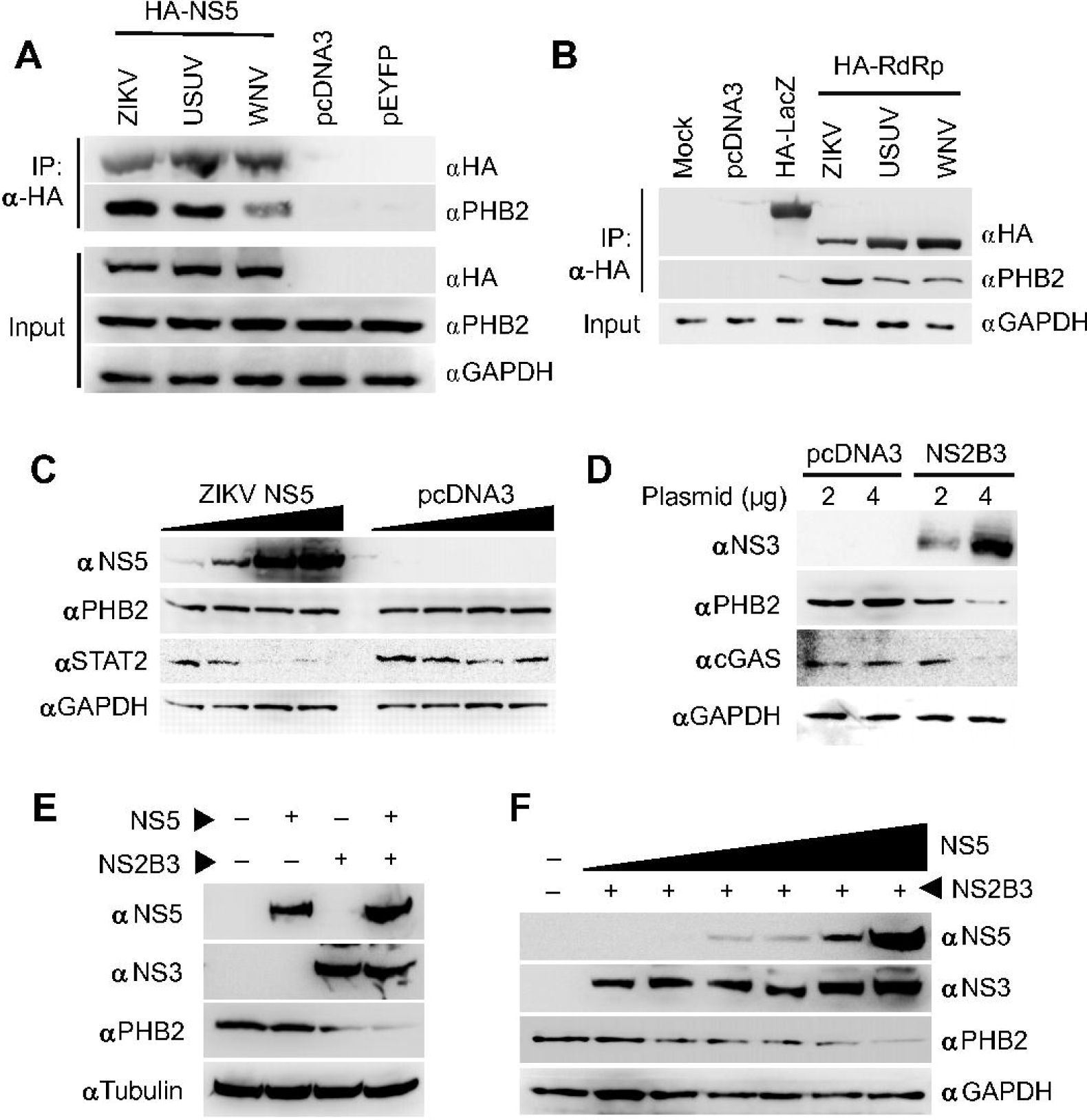
Ectopic co-expression of NS2B3 and NS5 mediates efficient PHB2 elimination. (A, B) PHB2 is co-immunoprecipitated with ZIKV, USUV and WNV NS5 proteins (A) or their RdRp domains (B). To this, 293T cells were transfected with constructs expressing HA-tagged versions of each NS5 (A), or the RdRp domain alone (B). As controls, there were included mock-transfected cells or constructs expressing EYFP (pEYFP-N1, shown as pEYFP), HA-tagged LacZ (pCruz HA-LacZ, shown as HA-LacZ) or an empty vector (pcDNA3). Cell lysates were incubated with anti-HA agarose beads binding to NS5 and RdRp, and the presence of PHB2 in the co-IPs was examined with a specific antibody. (C) ZIKV NS5 overexpression leads to reduced STAT2 but not PHB2 levels. 293T cells were transfected with increasing amounts (400, 800, 1600, 3200 ng) of either a construct expressing HA-tagged ZIKV NS5 or an empty vector (pcDNA3). (D) Ectopic expression of NS2B/NS3 precursor (NS2B3) leads to reduced PHB2 levels in 293T cells. To this, 5 x 10^5^ 293T cells were seeded on a 35-mm plate and transfected with either 2 or 4 μg of pZIKV-NS2B3 or an empty vector (pcDNA3). (E) PHB2 elimination by NS2B3 is enhanced in the presence of NS5. In this assay, 5 x 10^5^ 293T cells were seeded on a 35-mm plate and co-transfected with 2 μg of pZIKV-NS2B3 and 2 μg of pZIKV-HA-NS5 or an empty vector (pcDNA3). (F) Larger amounts of NS5 lead to reduced PHB2 levels in cells expressing a suboptimal amount of NS2B/NS3. Transfections were carried out upon 5 x 10^6^ 293T cells on 100-mm plates with 2 μg of pZIKV-NS2B3 and different amounts of pZIKV-HA-NS5 (gradually increasing from 0.5 to 6 μg).

Since NS5 is not sufficient for PHB2 degradation, we anticipated that other viral factors might be involved. In addition to viral polyprotein processing, NS2B/NS3 also participates in the proteolytic cleavage of different cellular proteins (60–62). We confirmed that the ectopic expression of NS2B/NS3 precursor (NS2B3) resulted in decreased PHB2 levels (Fig. 9D). To examine a possible synergistic activity of both viral proteins, we analysed the effect of NS5 coexpression in cells transfected with a suboptimal amount of pZIKV-NS2B3. When both NS2B/NS3confirming that, when both proteins are present, a lower amount of PHB2 is detected (Figs 9E and 9F).

## 6. DISCUSSION

With the aim advancing our knowledge on the complex virus-host networks controlling genome replication, we characterised the cellular proteins interacting with three different NS5s. As a result of this analysis, we have obtained evidence supporting that PHB2, a mitophagy receptor, is a restriction factor for OFV replication. Previous studies have established a relationship between the OFVs and mitophagy (63). While its activation is necessary for Japanese encephalitis virus replication in a hepatoma cell line (64), mitophagy is silenced during DENV infection (65). For ZIKV, mitophagy activation has been associated with both proviral and antiviral activities (63). Ponia and colleagues demonstrated that ZIKV supresses mitophagy in astrocytes, in a process where NS5 interferes with host protein Ajuba (38). By contrast, Lee and Shin found that mitophagy inhibition, through RNAi-mediated silencing of PINK1, repressed viral replication, suggesting that mitophagy is distinctly regulated by different OFVs and/or cellular contexts (63, 66).

In our study, PHB2 depletion resulted in increased viral titres, genomic RNA and protein translation, sustaining that this protein is a restriction factor (Fig. 5). In addition to PHB2, we have identified other mitochondrial proteins as interactors, such as phosphoglycerate mutase family member 5 (PGAM5), which also plays an important role during mitophagy (40, 67), several subunits of the mitochondrial ATP synthase (e.g. ATP5F1B), fumarate hydratase (FH) and sideroflexin-1 (SFXN1). These findings are suggestive of mitochondria dynamics playing a crucial role during infection, and of NS5 as a factor subverting some of the processes controlled by these organelles. Despite this previous connection between ZIKV NS5 and mitophagy, it is conceivable, however, that viral restriction is linked to other cellular processes controlled by PHB2. A tantalising possibility is that PHB2 antiviral activity is related to its nuclear role as an ESR repressor (68–70). Supporting this possibility, inhibition of the oestrogen-dependent signal by tamoxifen also affects orthoflaviviral replication (70). Additional studies on ESR and other transcription factors regulated by PHB2 are necessary to fully understand this antiviral activity.

PHB2 is transiently eliminated during infection in different human cell contexts (Fig. 7). Since both ZIKV and DENV NS5 proteins target STAT2 for degradation in a ubiquitin-dependent manner we initially considered this possibility for PHB2. In this process, orthoflaviviral NS5s act as a bridge between STAT2 and an E3 ubiquitin ligase (35, 37). We have recently documented that OFV NS5s also interact with the main E1 ubiquitin-activating enzyme UBA1 (32). However, we found that the ectopic expression of NS5 does not affect PHB2 levels, being required the viral protease NS2B/NS3 to its degradation (Fig. 9D). When NS2B/NS3 is expressed at suboptimal levels, NS5 is needed for PHB2 elimination, suggesting that NS5 operates as an adaptor between the viral protease and the host factor (Figs 9E and 9F). A physical interaction between NS5 and NS3 has been previously documented, and hence it is also expected to take place between NS5 and the NS2B/NS3 complex (71, 72). An attractive possibility is that NS5 could be acting as a protein bond between NS2B/NS3 and various host restriction factors, in addition to PHB2.

Previous studies have established a connection between PHB2 and MAVS antiviral signalling (44). Therefore, a forthcoming explanation would be that OFVs targeting PHB2 to degradation might help uncouple MAVS from IFN-I expression (44). Our data suggest however that PHB2 knockdown activates IFN-I transcription, with larger IFNB1 mRNA levels in either infected or poly I:C–stimulated cells (Fig. 8). Larger IFNB1 transcription levels observed in PHB2-silenced cells seems counterintuitive to the antiviral role proposed for this host protein (Fig. 8). This could be partially explained in the context of PHB2 fluctuating levels in the infected cells (Fig. 7). We hypothesise that transient PHB2 degradation at 48 hpi is necessary for efficient orthoflaviviral genome replication. However, subsequent PHB2 restoration to normal levels at 60 hpi is predicted to reactivate mitophagy, and thereby mitigate the ensuing inflammation and MAVS-mediated IFN-I responses (73–76).

Gene downregulation assays have identified other NS5-interacting host partners affecting orthoflaviviral replication. Larger virus titres were recovered in RSL1D1-depleted cells, also supporting an antiviral activity associated with this cellular factor (Fig. S6). RSL1D1 is an RNA-binding protein participating in multiple cellular processes, including antagonising senescence, proliferation and apoptosis, and acting as an autophagy suppressor in different types of cancer (77, 78). Conversely, MYOF and LGALS3BP downregulation led to significant decreases in both ZIKV and USUV titres, indicating that they may be contributing to critical steps of the life cycle (Fig. 4 and Fig. S4). MYOF is a multifunctional Ca2+ – binding protein which regulates vesicle trafficking and the cellular membrane repair (79, 80). Since OFVs assemble their replication sites on ER invaginations, requiring extensive membrane remodelling, membrane-repair activities might be relevant to their formation (81). WJ460, a promising anticancer drug targeting MYOF Ca2+ – binding C2D domain (79, 82), showed only a minor effect on virus titres, in contrast with DsiRNA-mediated silencing (Fig. 4). Recent studies suggest that MYOF might not be the primary host target of WJ460, and thus the activities observed for this drug not being totally associated with this host factor (83). LGALS3BP is a secreted glycoprotein that has been related to immune modulation, tumour progression, and extracellular matrix organization. Mice infected with WNV show upregulated LGALS3BP expression in the CNS (84). However, while KO mice for this gene exhibit reduced neuroinflammation and alterations in T cell populations, no significant differences in viral titres were observed (84). This evidence suggests that LGALS3BP is not as necessary for WNV replication, as it appears to be for ZIKV and USUV, according to our study.

In addition to advance our understanding on the biological processes that control the orthoflaviviral life cycle, we deem these data presented here open new avenues for their therapeutic control. Approaches based on stimulating PHB2 antiviral activity or, alternatively, molecules triggering mitophagy activation, which are expected to antagonise viral replication, could be considered in future studies. The identification of the interacting epitopes between NS5 and other cellular partners can be of use to the rational design of host-targeting antivirals.

## Supporting information

Legends for supplemental material

## 7. ACKNOWLEDGEMENTS

We thank José Joaquín Lorenzo for his technical assistance to this work.

