## Supplementary table 1 for "Mitophagy-related prohibitin 2 is an orthoflavivirus restriction factor targeted for degradation during infection"

**Table S1. DsiRNAs used in the study:**

| **Host gene** | **Reference number** | **Dicer-substrate siRNA duplex (DsiRNA)** |
|---|---|---|
| Negative control | 51-01-14-04 | CGUUAAUCGCGUAUAAUACGCGU**AT**/<br>AUACGCGUAUUAUACGCGAUUAACGAC |
| ATP5F1B | hs.Ri.ATP5B.13.1 | GGUCAAGAUGUACUGCUAUUUAU**TG**/<br>CAAUAAAUAGCAGUACAUCUUGACCUU |
| DDX21 | hs.Ri.DDX21.13.1 | GGAUAGGCAUGGGUACCUUAUCA**AG**/<br>CUUGAUAAGGUACCCAUGCCUAUCCAA |
| FH | hs.Ri.FH.13.1 | CAGGUUUAAAUACUAGAAUUGGC**TT**/<br>AAGCCAAUUCUAGUAUUUAAACCUGUA |
| HSD17B12 | hs.Ri.HSD17B12.13.1 | ACCAAGAAGAACUAAGCAUUGAU**AA**/<br>5’UUAUCAAUGCUUAGUUCUUCUUGGUUU |
| LGALS3BP | hs.Ri.LGALS3BP.13.1 | AGUCACAACUGGUCUAUCAGUCC**AG**/<br>CUGGACUGAUAGACCAGUUGUGACUUC |
| MYOF | hs.Ri.MYOF.13.1 | GCUCUAAAAUCAGGGAUACAAGG**TA**/<br>UACCUUGUAUCCCUGAUUUUAGAGCUU |
| PGAM5 | hs.Ri.PGAM5.13.1 | CUUGGGGUUGAAGUUUAAUAAAA**TC**/<br>GAUUUUAUUAAACUUCAACCCCAAGCU |
| PHB2 | hs.Ri.PHB2.13.1 | GAAAUGAGCCUAGUCACCAAGAA**CT**/<br>AGUUCUUGGUGACUAGGCUCAUUUCUU |
| RPL21 | N/A | AUAAGAAAGGUGAUAUUGUAGAC**AT**/<br>GAUAUUCUUUCCACUAUAACAUCUGUA |
| RPL24 | hs.Ri.RPL24.13.1 | AGUUGGUGGAAAACGCUAAACUG**GC**/<br>GCCAGUUUAGCGUUUUCCACCAACUCG |
| RPL34 | hs.Ri.RPL34.13.1 | GUAAUAGAAUUGUUUACCUUUAU**AC**/<br>GUAUAAAGGUAAACAAUUCUAUUACCA |
| RSL1D1 | hs.Ri.RSL1D1.13.1 | AAGCAUGGAAUUAAAACCGUUUC**TC**/<br>GAGAAACGGUUUUAAUUCCAUGCUUGU |
| SMARCA5 | hs.Ri.SMARCA5.13.1 | AUCUGUUUACUGAGCUAGAAACA**TA**/<br>UAUGUUUCUAGCUCAGUAAACAGAUGA |
| SFXN1 | hs.Ri.SFXN1.13.1 | GGACGAGCCAAUCAUUUCUUCAC**TG**/<br>CAGUGAAGAAAUGAUUGGCUCGUCCAA |
