## Supplementary figures for "Mitophagy-related prohibitin 2 is an orthoflavivirus restriction factor targeted for degradation during infection"

**Fig. S1**

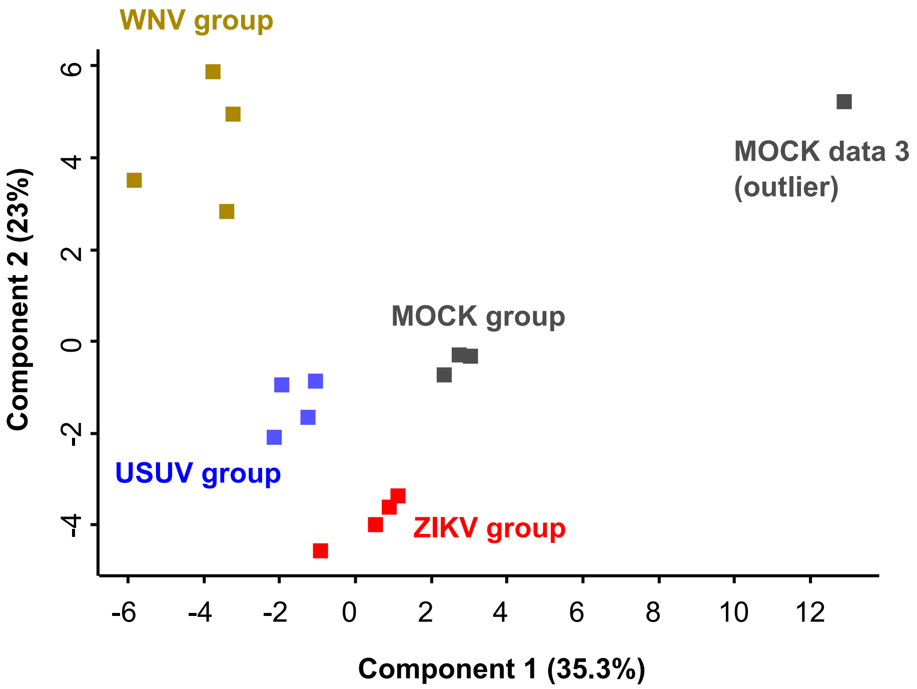
WNV group
MOCK data 3
(outlier)
MOCK group
USUV group
ZIKV group
Component 2 (23%)
Component 1 (35.3%)
6
4
2
0
-2
-4
-6
-4
-2
0
2
4
6
8
10
12

### Fig. S2

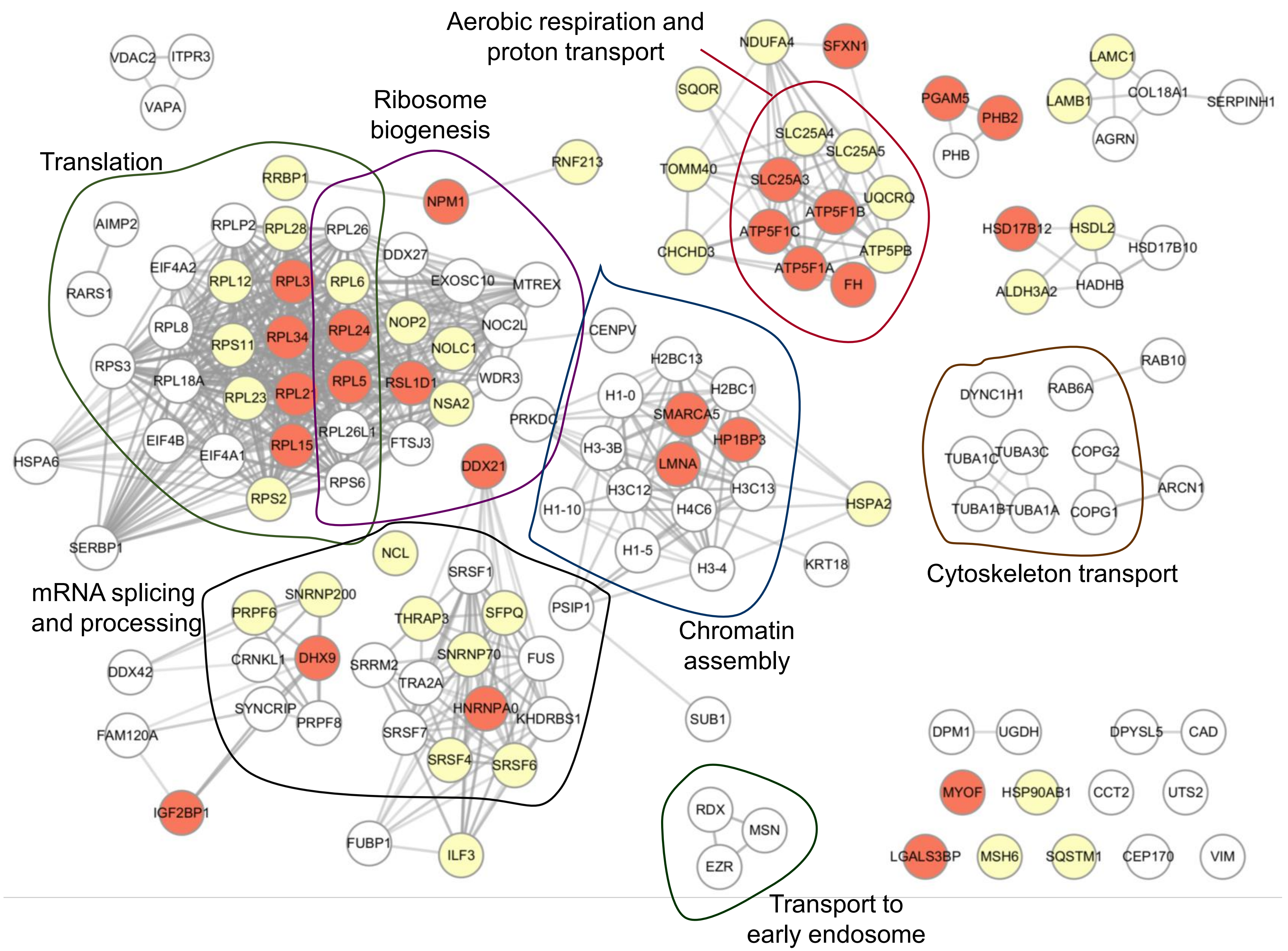

Aerobic respiration and proton transport
Ribosome biogenesis
Translation
Chromatin assembly
Cytoskeleton transport
mRNA splicing and processing
Transport to early endosome
VDAC2
ITPR3
VAPA
NDUFA4
SFXN1
SQOR
LAMC1
PGAM5
PHB2
LAMB1
COL18A1
SERPINH1
AGRN
PHB
SLC25A4
SLC25A5
TOMM40
SLC25A3
UQCRQ
ATP5F1B
ATP5F1C
CHCHD3
ATP5PB
ATP5F1A
FH
HSD17B12
HSDL2
HSD17B10
ALDH3A2
HADHB
RNF213
RRBP1
NPM1
AIMP2
RPLP2
RPL28
RPL26
DDX27
EIF4A2
RPL12
RPL3
RPL6
EXOSC10
MTREX
RARS1
RPL8
RPS11
RPL34
RPL24
NOP2
NOC2L
NOLC1
CENPV
RPS3
RPL18A
RPL23
RPL21
RPL5
RSL1D1
WDR3
NSA2
PRKDC
RPL26L1
EIF4B
EIF4A1
RPL15
FTSJ3
HSPA6
RPS6
DDX21
RPS2
SERBP1
H2BC13
H2BC1
H1-0
SMARCA5
HP1BP3
H3-3B
LMNA
H3C12
H3C13
H1-10
H4C6
H1-5
H3-4
HSPA2
KRT18
RAB10
DYNC1H1
RAB6A
TUBA1C
TUBA3C
COPG2
ARCN1
TUBA1B
TUBA1A
COPG1
NCL
SRSF1
SNRNP200
PRPF6
THRAP3
SFPQ
PSIP1
CRNKL1
DHX9
SRRM2
SNRNP70
FUS
DDX42
TRA2A
SYNCRIP
PRPF8
HNRNPA0
KHDRBS1
FAM120A
SRSF7
SRSF4
SRSF6
SUB1
IGF2BP1
FUBP1
ILF3
RDX
MSN
EZR
DPM1
UGDH
DPYSL5
CAD
MYOF
HSP90AB1
CCT2
UTS2
LGALS3BP
MSH6
SQSTM1
CEP170
VIM

**Fig. S3**

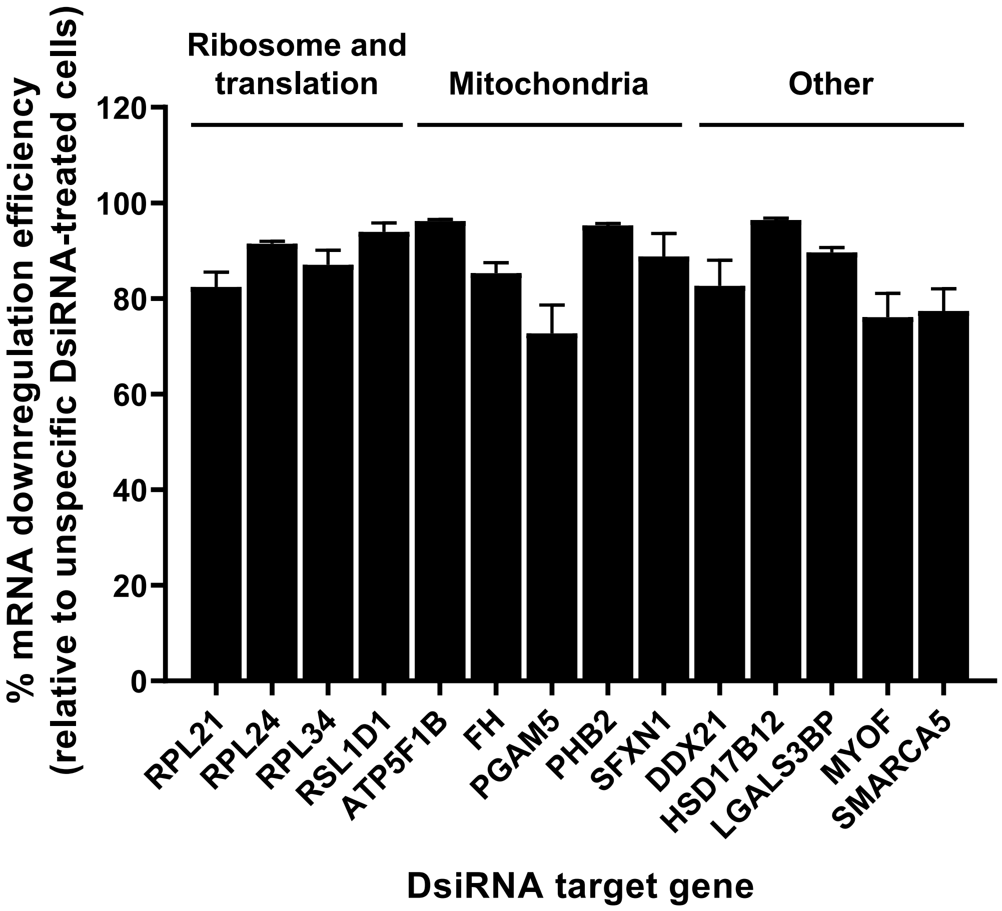
Ribosome and translation
Mitochondria
Other
% mRNA downregulation efficiency
(relative to unspecific DsiRNA-treated cells)
120
100
80
60
40
20
0
RPL21
RPL24
RPL34
RSL1D1
ATP5F1B
FH
PGAM5
PHB2
SFXN1
DDX21
HSD17B12
LGALS3BP
MYOF
SMARCA5
DsiRNA target gene

### Fig. S4

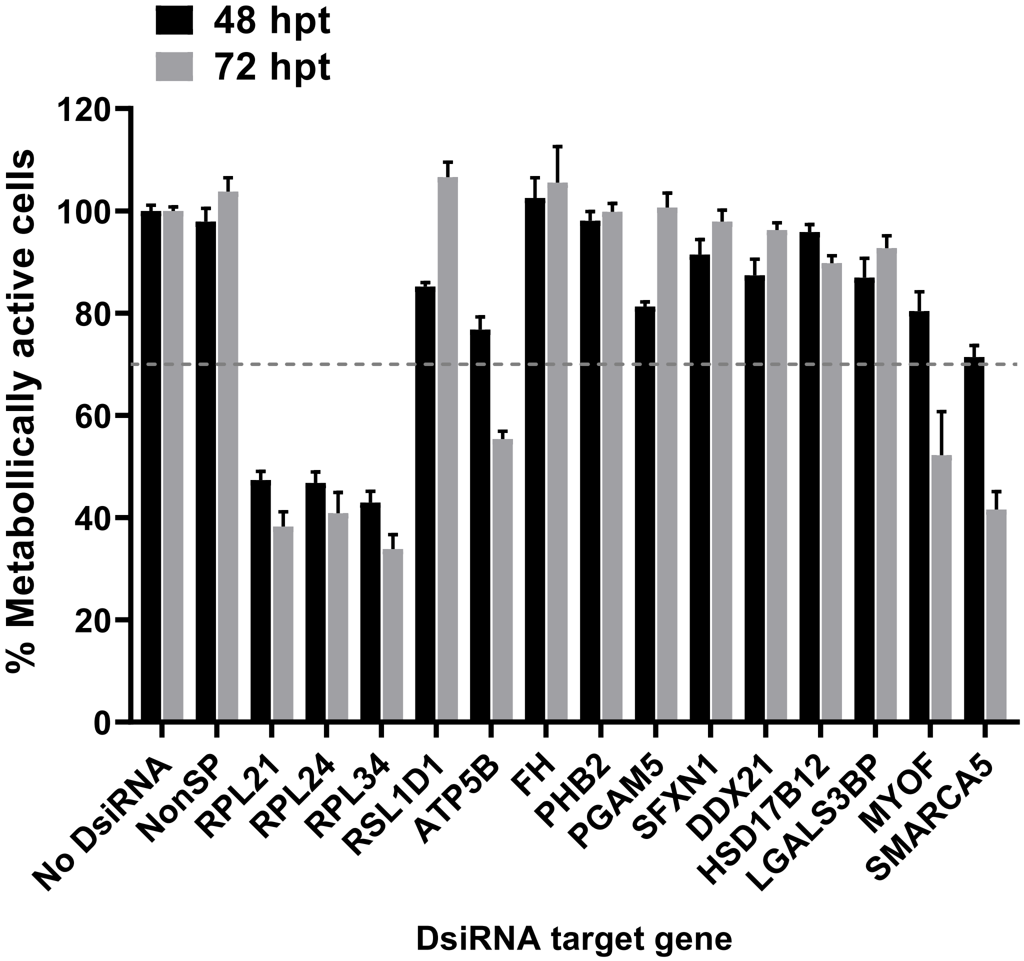
48 hpt
72 hpt
% Metabollically active cells
0
20
40
60
80
100
120
No DsiRNA
NonSP
RPL21
RPL24
RPL34
RSL1D1
ATP5B
FH
PHB2
PGAM5
SFXN1
DDX21
HSD17B12
LGALS3BP
MYOF
SMARCA5
DsiRNA target gene

**Fig. S5**

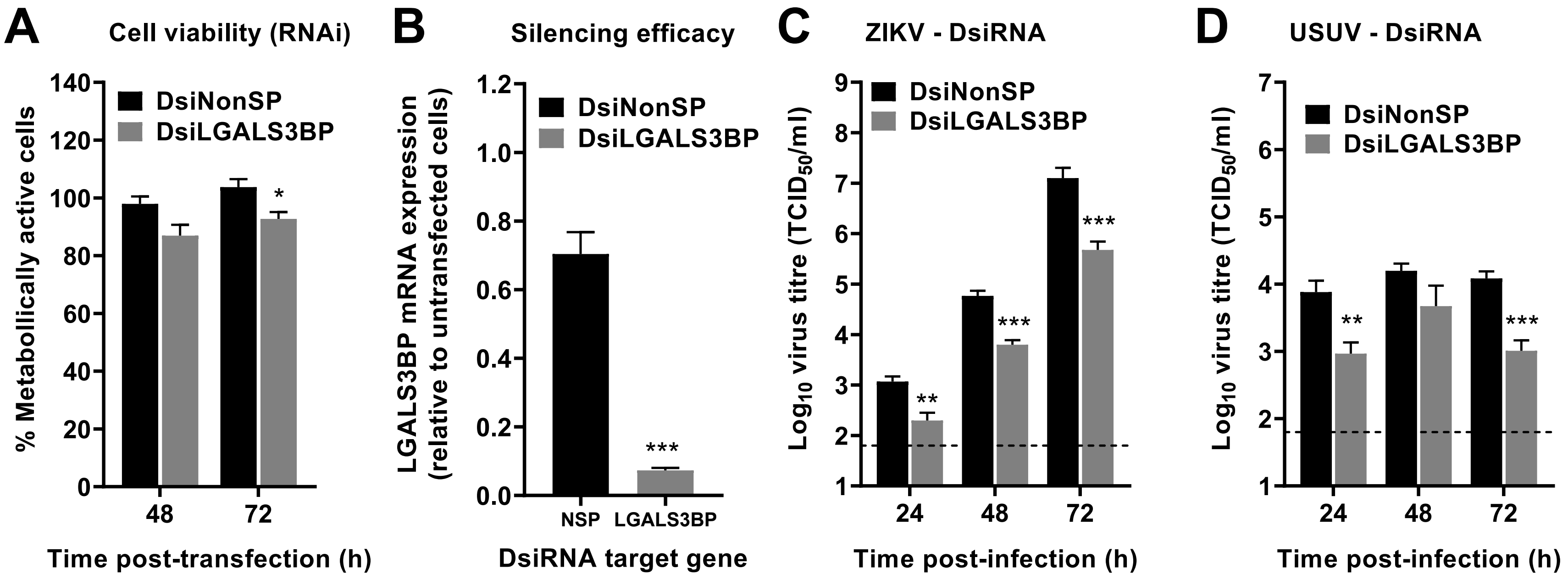
A
Cell viability (RNAi)
DsiNonSP
DsiLGALS3BP
% Metabolically active cells
140
120
100
80
60
40
20
0
*
48
72
Time post-transfection (h)
B
Silencing efficacy
LGALS3BP mRNA expression (relative to untransfected cells)
1.2
1.0
0.8
0.6
0.4
0.2
0.0
***
NSP
LGALS3BP
DsiRNA target gene
C
ZIKV - DsiRNA
Log10 virus titre (TCID50/ml)
9
8
7
6
5
4
3
2
1
**
***
***
24
48
72
Time post-infection (h)
D
USUV - DsiRNA
7
6
5
4
3
2
1
**
***
24
48
72
Time post-infection (h)

**Fig. S6**

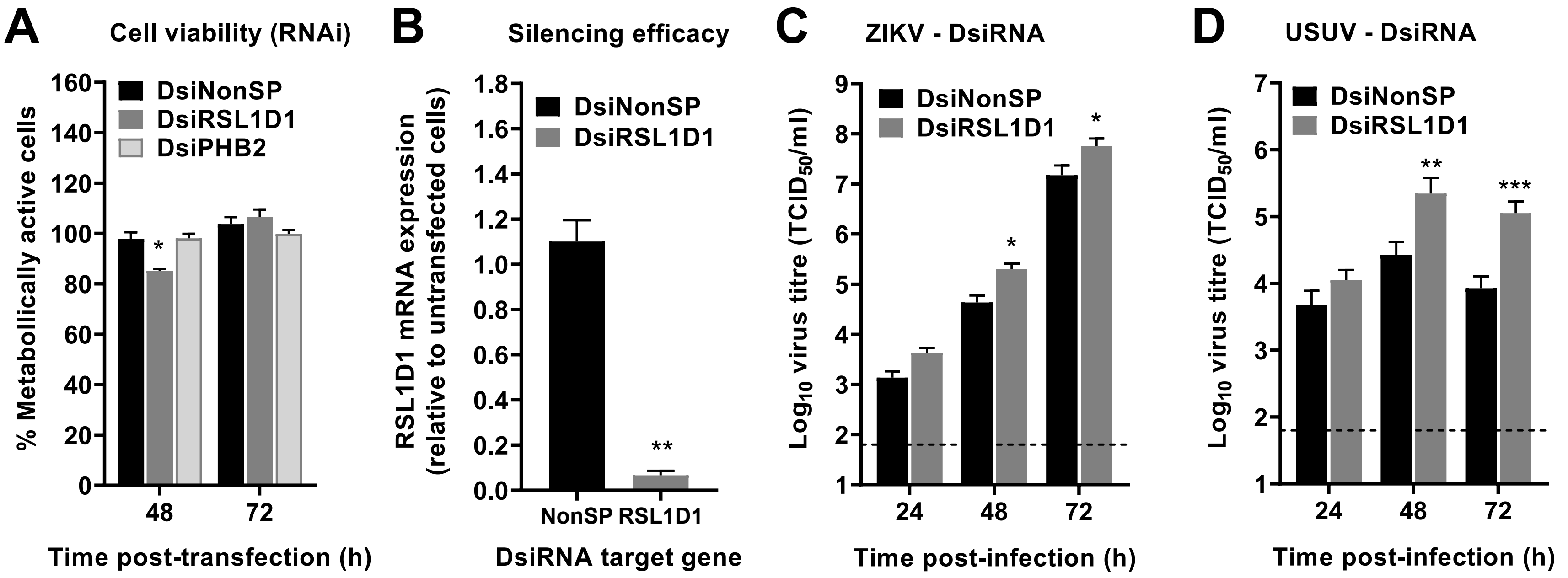
A
Cell viability (RNAi)
DsiNonSP
DsiRSL1D1
DsiPHB2
% Metabollically active cells
160
140
120
100
80
60
40
20
0
*
48
72
Time post-transfection (h)
B
Silencing efficacy
DsiNonSP
DsiRSL1D1
RSL1D1 mRNA expression (relative to untransfected cells)
1.8
1.6
1.4
1.2
1.0
0.8
0.6
0.4
0.2
0.0
**
NonSP
RSL1D1
DsiRNA target gene
C
ZIKV - DsiRNA
DsiNonSP
DsiRSL1D1
Log10 virus titre (TCID50/ml)
9
8
7
6
5
4
3
2
1
*
*
24
48
72
Time post-infection (h)
D
USUV - DsiRNA
DsiNonSP
DsiRSL1D1
Log10 virus titre (TCID50/ml)
7
6
5
4
3
2
1
**
***
24
48
72
Time post-infection (h)

Fig. S7

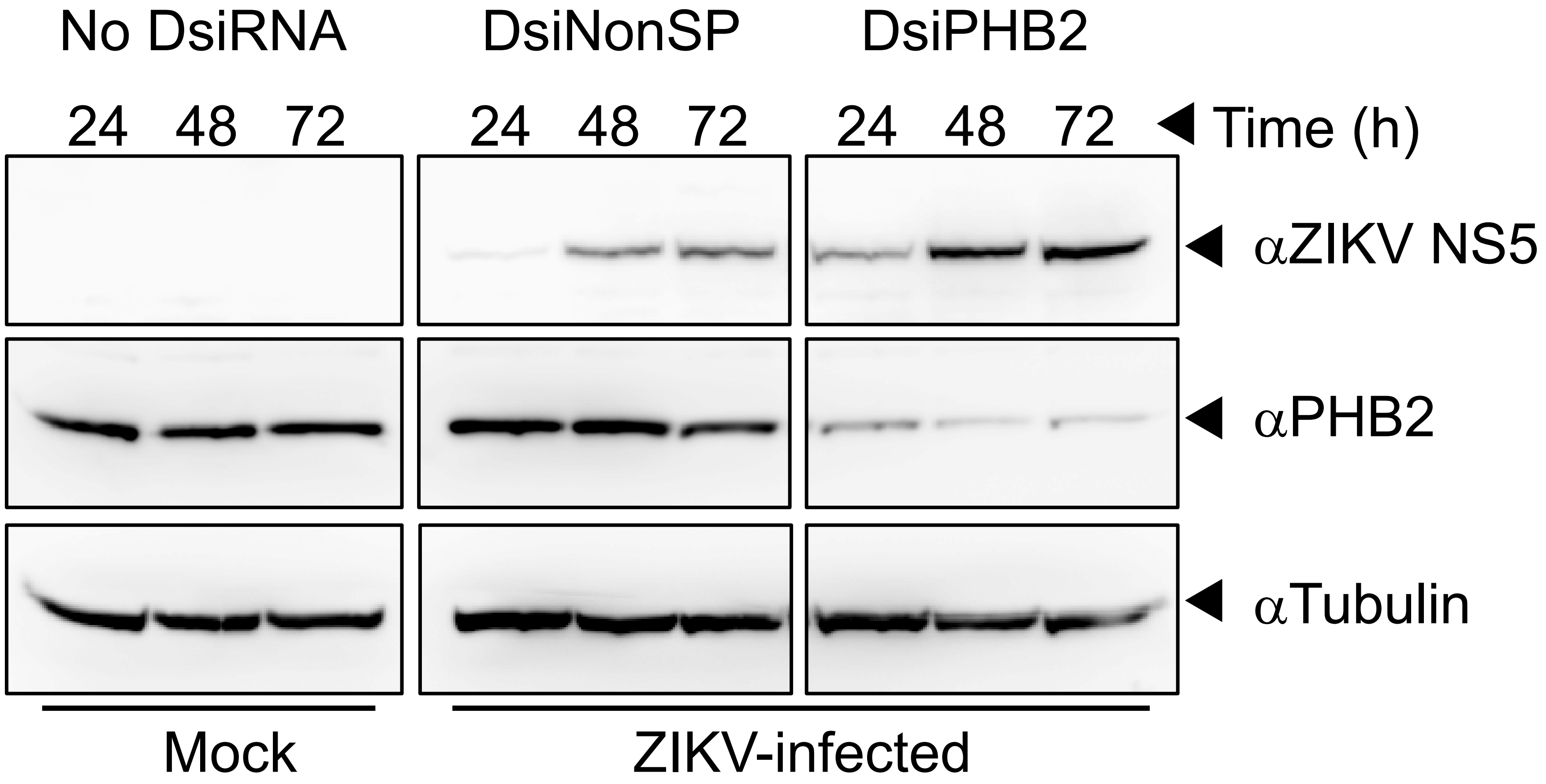
No DsiRNA
DsiNonSP
DsiPHB2
24 48 72
24 48 72
24 48 72
Time (h)
αZIKV NS5
αPHB2
αTubulin
Mock
ZIKV-infected

Fig. S8

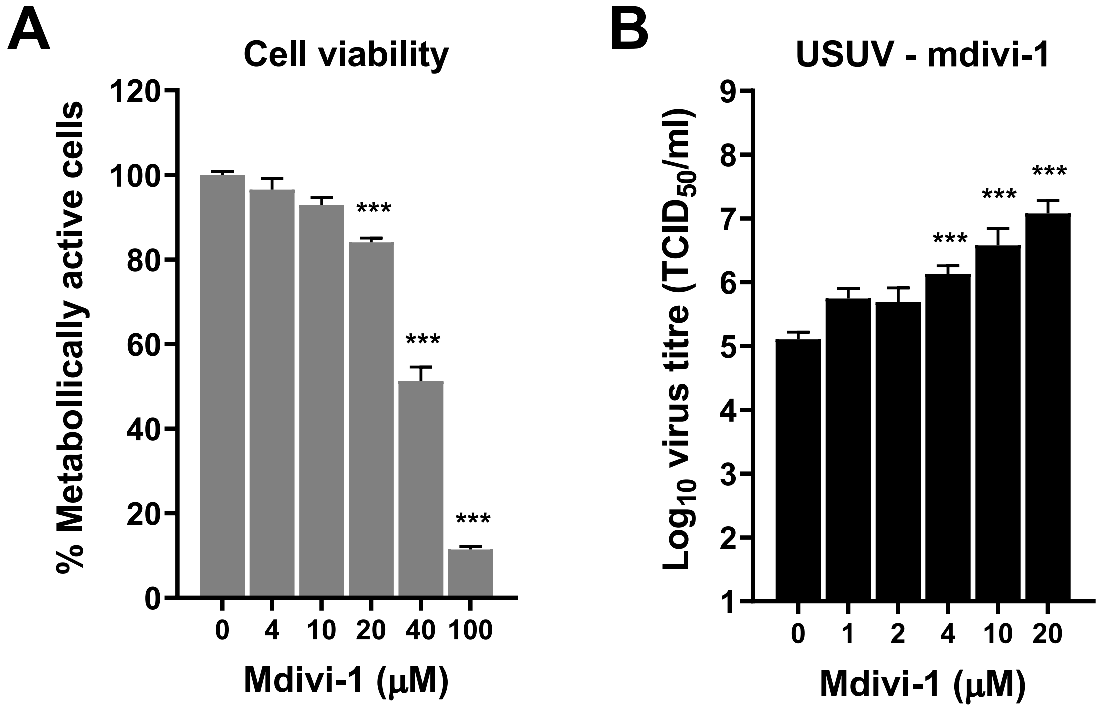
A
Cell viability
% Metabollically active cells
0
20
40
60
80
100
120
***
***
***
0 4 10 20 40 100
Mdivi-1 (µM)
B
USUV - mdivi-1
Log10 virus titre (TCID50/ml)
1
2
3
4
5
6
7
8
9
***
***
***
0 1 2 4 10 20
Mdivi-1 (µM)

**Fig. S9**

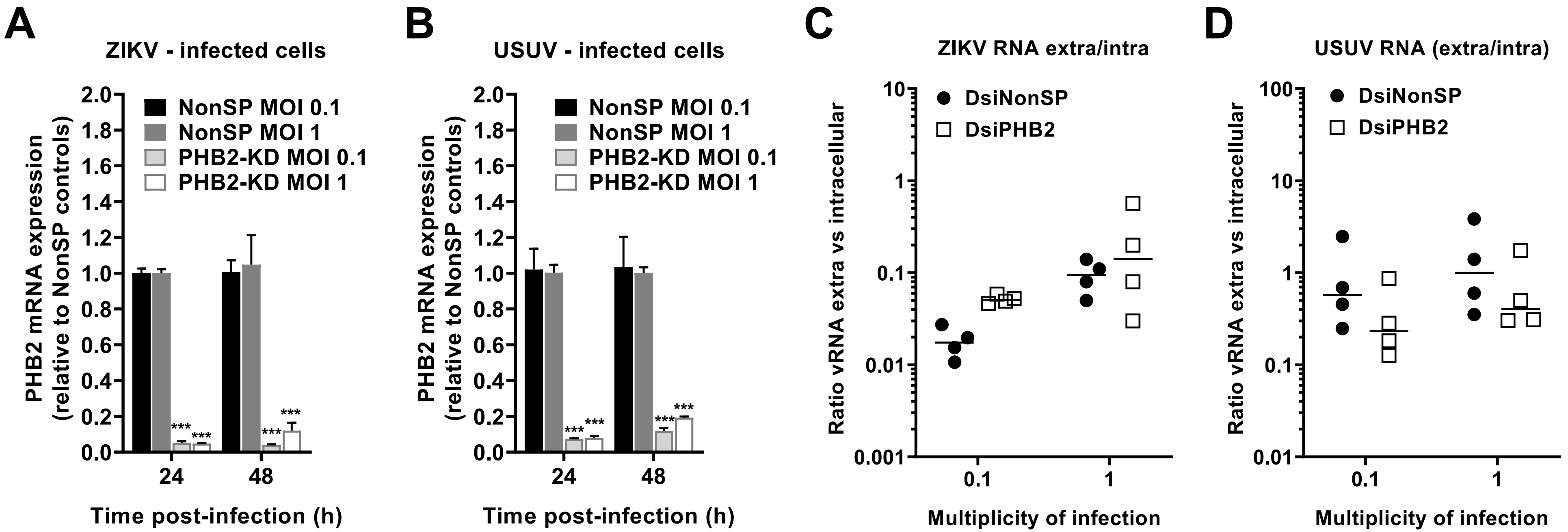
A
ZIKV - infected cells
PHB2 mRNA expression (relative to NonSP controls)
NonSP MOI 0.1
NonSP MOI 1
PHB2-KD MOI 0.1
PHB2-KD MOI 1
Time post-infection (h)
B
USUV - infected cells
C
ZIKV RNA extra/intra
Ratio vRNA extra vs intracellular
DsiNonSP
DsiPHB2
Multiplicity of infection
D
USUV RNA (extra/intra)

### Fig. S10

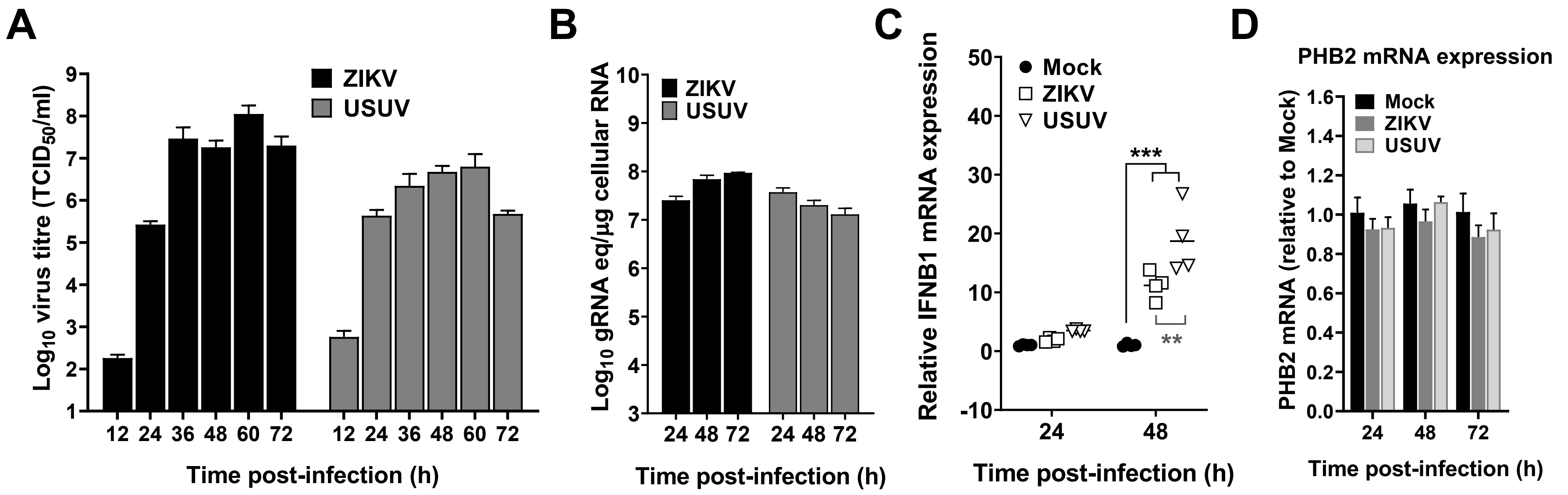
A
ZIKV
USUV
Log10 virus titre (TCID50/ml)
9
8
7
6
5
4
3
2
1
12 24 36 48 60 72 12 24 36 48 60 72
Time post-infection (h)
B
ZIKV
USUV
Log10 gRNA eq/μg cellular RNA
10
9
8
7
6
5
4
3
24 48 72 24 48 72
Time post-infection (h)
C
Mock
ZIKV
USUV
Relative IFNB1 mRNA expression
50
40
30
20
10
0
-10
***
**
24
48
Time post-infection (h)
D
PHB2 mRNA expression
Mock
ZIKV
USUV
PHB2 mRNA (relative to Mock)
1.6
1.4
1.2
1.0
0.8
0.6
0.4
0.2
0.0
24
48
72
Time post-infection (h)

**Fig. S11**

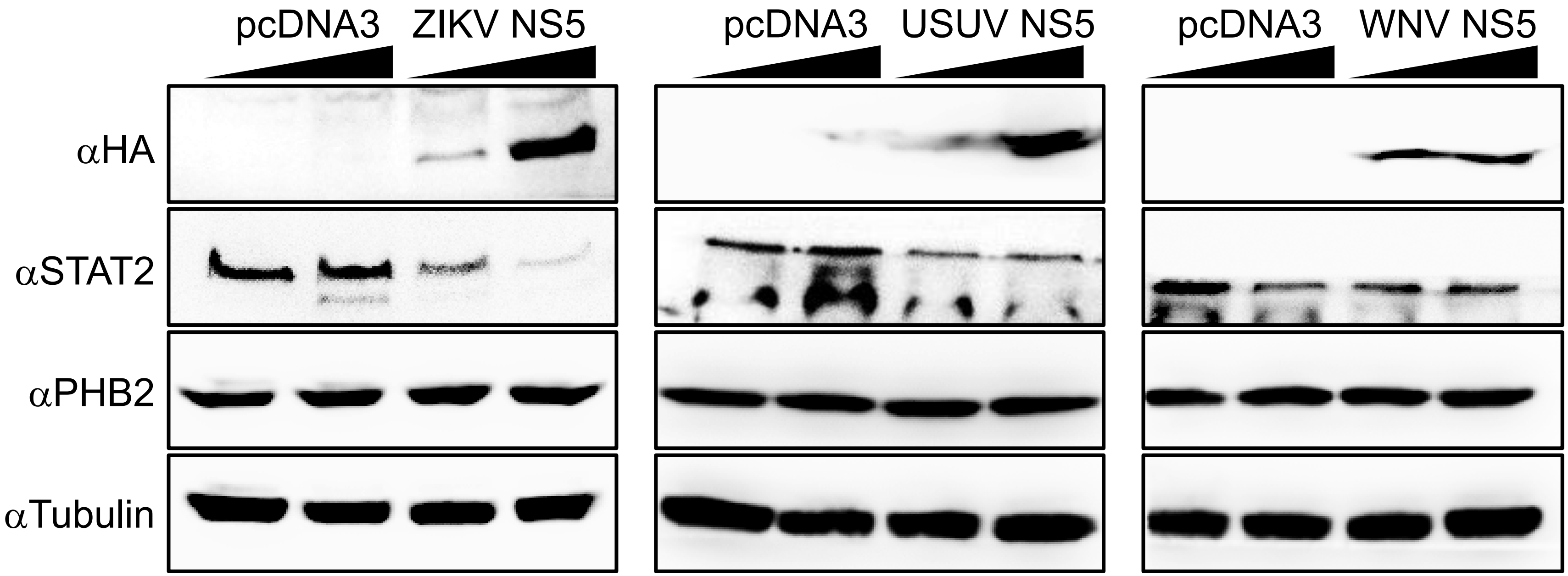
pcDNA3
ZIKV NS5
pcDNA3
USUV NS5
pcDNA3
WNV NS5
αHA
αSTAT2
αPHB2
αTubulin
