## Supplementary material for "Mitophagy-related prohibitin 2 is an orthoflavivirus restriction factor targeted for degradation during infection": Legends for supplemental material

**Legends of Supplemental material for manuscript:**

Affiliations:

### 1. SUPPLEMENTARY FIGURE LEGENDS

**Fig. S1. Quality control of the proteomic datasets.** Principal component analysis (PCA) separates each coimmunoprecipitation (co-IP) dataset (n=4) from each other (ZIKV, USUV and WNV) and from control co-IP in mock-transfected cells. PCA also shows reproducible clustering of biological replicates, except for one control experiment (outlier) that has been removed for further analyses.

**Fig. S2. Orthoflavivirus NS5 interaction network.** Quantitative proteomics was used as described in the text to identify host factors interacting with OFV NS5 proteins. Briefly, host protein interactors were pulled down from cell lysates obtained after transfection of HA-tagged versions of ZIKV, USUV or WNV NS5 proteins and subsequent immunoprecipitation. A total of 141 proteins were identified after combining all three datasets. A protein-protein interaction network for these factors was generated using the STRING database (1). Data visualisation was performed using Cystoscape (2), and clustering to identify densely connected nodes was then applied to the network with the MCL algorithm. Those factors identified in two datasets were represented in pale yellow. Those factors found in three datasets were shown either in coral (top 50 enriched factors in all three datasets) or in light orange (remaining host factors). The biological process term (GO-BP) better fitting each cluster is given.

**Fig. S3. Relative knockdown efficacy of host-specific DsiRNAs in A549 cells.** RT-qPCR analysis at 48 h post-transfection of host gene expression in A549 cells lipofected with a host-specific DsiRNA and normalised to GAPDH mRNA levels. Values obtained are the average of ≥ 4 independent biological replicas. Knockdown efficiency was calculated by RT-qPCR and the $2^{-\Delta\Delta Ct}$ method, using as a reference a cell culture transfected with a non-human target DsiRNA.

**Fig. S4. Effect of host factor depletion on cell viability.** Relative number of metabolically active cells after specific host gene downregulation compared to mock-transfected cells. Cells were lipofected with either a host factor-specific DsiRNA or an unspecific DsiRNA (NonSP). The relative number of viable cells (metabolically active) is calculated with the *CellTiter-Blue Cell Viability Assay* kit at 48 h (black) and 72 h post-transfection (grey). A dashed line is included to indicate those viability values above 70%.

**Fig. S5. Reduced virus titres in LGALS3BP-depleted cells.** (A) Relative amount of metabolically active cells after transfection with a specific (DsiLGALS3BP) or an unspecific DsiRNA (DsiNonSP). All values are relative to untransfected cells (not represented in the graph). (B) RT-qPCR analysis of LGALS3BP expression in A549 cells transfected with either a host-specific (DsiLGALS3BP) or a non-human target DsiRNA (DsiNonSP). LGALS3BP mRNA expression was normalised to GAPDH mRNA levels. Values obtained are the average of at least 4 independent biological replicas ($n \geq 4$), and relative to untransfected cells carried out in parallel. (C, D) Virus infectivity was quantitated by $TCID_{50}$ assays in samples collected at different time points from cells previously transfected with 5 pmol of each DsiRNA, as indicated in A and B. Twenty-four hours after transfection, cells were infected with ZIKV (C) or USUV (D) at an MOI of 0.1 $TCID_{50}$/cell. Each value in the graph is the average of ≥ 4 independent biological replicas. The statistical significance was calculated by Two-way ANOVA using Sidak's multiple comparisons (A, C, D) or *t* (B) tests (*, $p<0.05$; **, $p<0.01$; ***, $p < 0.001$). Bars are representing the standard error of the mean (SEM) for each data point.

**Fig. S6. Increased virus titres in RSL1D1-depleted cells.** (A) Relative amount of metabolically active cells after transfection with a specific (DsiRSL1D1 or DsiPHB2) or an unspecific DsiRNA (DsiNonSP). Cell viability is determined at 48 and 72 h post-transfection as suggested in the Methods section, and relative to untransfected cells. (B) RT-qPCR analysis of RSL1D1 expression in A549 cells transfected with either an RSL1D1-targeting (DsiRSL1D1) or a non-human target DsiRNA (DsiNonSP). RSL1D1 mRNA expression was normalised to GAPDH mRNA levels. Values obtained are the average of 4 independent biological replicas (n=4), and relative to untransfected cells carried out in parallel. (C, D) Virus infectivity was examined in samples collected from the supernatants of cells previously transfected with each DsiRNA. Twenty-four hours after DsiRNA transfection, cells were infected with ZIKV (C) or USUV (D) at an MOI of 0.1 $TCID_{50}$/cell. Each value in the graph is the average of ≥ 4 independent biological replicas. (A – D) The statistical significance was calculated by Two-way ANOVA and Sidak's multiple comparison test (A, C, D) or a *t* test (B) (*, $p<0.05$; **, $p<0.01$; ***, $p < 0.001$). Bars are representing the standard error of the mean (SEM) for each data point.

**Fig. S7. Increased ZIKV NS5 protein levels in PHB2-depleted cells.** Representative Western blot of NS5 and PHB2 in ZIKV-infected cells previously transfected with either a specific (DsiPHB2) or a non-human specific (DsiNonSP) DsiRNA. At 24 h post-transfection, ZIKV was applied to cells at an MOI of 1 $TCID_{50}$/cell. Cell lysates were collected at different time points and analysed by Western blot. As a loading reference, an anti-tubulin antibody was used. Untransfected mock-infected cells were included as a reference control.

**Fig. S8. Extracellular to intracellular viral RNA levels in PHB2-downregulated cells**. (A – D) A549 cells were transfected with either a non-human targeting (NonSP) or a PHB2-targeting DsiRNA (PHB2). At 24 hours post-transfection, cells were infected with either ZIKV (A, C) or USUV (B, D) at an MOI of 0.1 or 1 $TCID_{50}$/cell. (A, B) RT-qPCR analysis of PHB2 expression in A549 cells transfected with either a PHB2-targeting (DsiPHB2) or a non-human specific (DsiNonSP) DsiRNA. PHB2 mRNA expression was normalised to GAPDH mRNA levels. Values are the average of 4 independent biological replicas (n=4), and relative to controls (NonSP) for each virus, MOI and time point represented. (C, D) Ratios of extracellular to intracellular viral RNA in PHB2-depleted cells at 48 h after infection with ZIKV (C) or USUV (D). The statistical significance was calculated in each graph by Two-way ANOVA and Sidak's multiple comparisons test. Bars are representing the standard error of the mean (SEM) for each data point.

**Fig. S9. Larger USUV titres in the presence of mitophagy inhibitor mdivi-1**. (A) Relative number of metabolically active A549 cells after 48 hours in the presence of increasing concentrations of mdivi-1. (B) USUV titres in the supernatant of infected cells after treatment with increasing concentrations of mdivi-1 during 48 h. Cells were previously inoculated with USUV at an MOI of 0.1 $TCID_{50}$/cell.

**Fig. S10. ZIKV and USUV replication kinetics and their effect on PHB2 and IFNB1 mRNA expression.** (A) Virus titre kinetics in the supernatant of A549 cells infected with either ZIKV (black) or USUV (grey). Bars are representing the standard error of the mean (SEM) for each data point. (B) Viral genome equivalents in cells infected with ZIKV (black) or USUV (grey). (C) Relative IFNB1 mRNA levels in cells infected with ZIKV or USUV. (D) RT-qPCR analysis of PHB2 mRNA expression in A549 cells infected with ZIKV or USUV. PHB2 was normalised to GAPDH mRNA levels and expressed as relative values to mock-infected cells carried out in parallel. Values obtained are the average of ≥ 4 independent biological replicas. The statistical significance was calculated by Two-way ANOVA and Sidak's multiple comparison test (A, C, D) or a *t* test (B) (*, $p<0.05$; **, $p<0.01$; ***, $p < 0.001$).

**Fig. S11. ZIKV NS5 overexpression leads to reduced STAT2 but not PHB2 protein levels**. Either 1200 or 2400 ng of constructs expressing ZIKV (right panel), USUV (middle) or WNV (left) NS5 or an empty vector (pcDNA3) are transfected in 293T cells. The effect of NS5 on the amount of STAT2 and PHB2 is examined by Western blot. Tubulin is used as a loading control.

### 2. SUPPLEMENTARY TABLE LEGENDS

**Table S1. DsiRNA molecules used in the study**. Nucleotide sequences of DsiRNA duplexes used for specific host gene silencing. Each DsiRNA duplex is formed by a 25-nt sense (above) and 27-nt antisense (below) RNA strand. The last two nucleotides found in each sense strand are deoxyribonucleotides (bold typed) while the remaining are ribonucleotides. All duplexes are commercially available at IDT, except the DsiRNA for RPL21 that consequently was designed by us. Its synthesis was then requested to the same company (IDT).

**Table S2. Summary of ZIKV NS5-host protein interaction networks**. A list of all factors identified in the analysis, using the *Perseus* software, is included. The top 50 significantly-enriched host interactors are shaded in pink. In grey, there have been highlighted the detection of HA-tagged ZIKV NS5 (correctly assigned) and USUV NS5.

**Table. S3. Summary of USUV NS5-host protein interaction networks**. A list of all factors identified in the analysis, using *Perseus* as abovementioned, is included. The top 50 significantly-enriched host interactors are shaded in pink. In grey, there have been highlighted the detection of HA-tagged USUV (correctly assigned) and WNV NS5 proteins.

**Table S4. Summary of WNV NS5-host protein interaction networks**. A list of all factors identified in the analysis, using *Perseus* as abovementioned, is included. The top 50 significantly-enriched host interactors are shaded in pink. WNV NS5 protein identification is highlighted in grey.

**Table S5. Summary of Biological Process (BP) enrichment analysis of orthoflavivirus NS5-host protein interactions**. To this analysis, we used all the host factors identified in Tables S2 – S4, and the g:GOSt application available at the g:Profiler website (3).

**Table S6. Summary of Cellular Compartments (CC) enrichment analysis of orthoflavivirus NS5-host protein interactions**. To this analysis, we used all the host factors identified in Tables S2 – S4, and the g:GOSt application available at the g:Profiler website (3).

### 3. REFERENCES (SUPPLEMENTAL DATA)


1. Szklarczyk D, Kirsch R, Koutrouli M, Nastou K, Mehryary F, Hachilif R, Gable AL, Fang T, Doncheva NT, Pyysalo S, Bork P, Jensen LJ, von Mering C. 2023. The STRING database in 2023: protein–protein association networks and functional enrichment analyses for any sequenced genome of interest. Nucleic Acids Res 51:D638–D646.

2. Shannon P, Markiel A, Ozier O, Baliga NS, Wang JT, Ramage D, Amin N, Schwikowski B, Ideker T. 2003. Cytoscape: A Software Environment for Integrated Models of Biomolecular Interaction Networks. Genome Res 13:2498–2504.

3. Kolberg L, Raudvere U, Kuzmin I, Adler P, Vilo J, Peterson H. 2023. g:Profiler—interoperable web service for functional enrichment analysis and gene identifier mapping (2023 update). Nucleic Acids Res 51:W207–W212.
